# Electrostatic control of chromatin compaction safeguards against apoptotic DNA release

**DOI:** 10.64898/2026.02.23.707452

**Authors:** Maximilian F. D. Spicer, Sanne Wijma, Nikki Schütte, Jan Huertas, M. Julia Maristany, José Ignacio Pérez López, Lifeng Chen, Mustafa Alaabo, Michael K. Rosen, Rosana Collepardo-Guevara, Daniel W. Gerlich

## Abstract

Apoptosis involves extensive intracellular reorganisation to facilitate the clearance of dying cells. A key step in this process is the destruction of the genome through fragmentation by caspase-activated endonuclease (CAD). Rather than dispersing after CAD-mediated cleavage, DNA fragments are compacted into a dense chromatin compartment. However, the underlying mechanism and biological relevance of this compaction remain unknown. Here we show that global deacetylation of histone tails promotes chromatin compaction during apoptosis, preventing DNA release into apoptotic extracellular vesicles. Using synthetic effectors that modulate nucleosome electrostatics independently of histone modifications, we demonstrate that electrostatic attraction alone is sufficient to compact and sequester fragmented chromatin. These findings reveal a mechanism by which global reprogramming of histone modifications coordinates fragmentation of the genome with its physical sequestration during apoptosis. Furthermore, our synthetic approach provides a tool to probe the role of physical forces in genome organisation across diverse biological contexts.

---

Programmed cell death by apoptosis is essential for the development and homeostasis of multicellular organisms, enabling the orderly elimination of superfluous or damaged cells without compromising tissue integrity^1–8^. To facilitate efficient clearance, apoptotic cells undergo a coordinated disassembly programme that reorganises intracellular structures into discrete compartments suitable for phagocytic uptake^5,6,8,9^. A hallmark of this disassembly process is the formation of apoptotic extracellular vesicles (ApoEVs), which disseminate cellular components into the surrounding tissue environment and can engage immune cells to modulate downstream responses^5,10,11^. While ApoEVs generally incorporate proteins, lipids, and RNA, nuclear DNA fragments generated by CAD^12–15^ are strikingly absent from most vesicles^16,17^. This selective DNA exclusion is thought to limit inflammatory responses, given the immunostimulatory potential of extracellular DNA^18,19^ and the involvement of DNA-containing extracellular vesicles in autoimmune disorders such as lupus erythematosus^20,21^. However, the mechanisms preventing DNA entry into vesicles during apoptosis remain unclear.

Chromatin compaction is a conserved morphological feature of apoptosis^10,22^, raising the possibility that compaction functions as a mediator of DNA sequestration. However, because the mechanism of apoptotic chromatin compaction remains undefined, its role in DNA sequestration has not yet been tested. Here, we demonstrate that histone deacetylation and changes in chromatin electrostatics drive chromatin compaction during apoptosis, thereby enabling genome fragmentation without the release of DNA beyond the cell corpse.

## Histone deacetylation drives apoptotic chromatin compaction

Genome-wide chromatin compaction is a hallmark of apoptosis, yet the molecular mechanism that drives this transformation remains unresolved. Given that mitotic chromosome compaction relies on global histone deacetylation^23–25^, we considered whether a similar mechanism might underlie chromatin reorganisation during apoptosis. Although histone deacetylation has been observed in apoptotic cells^26–29^, its functional contribution to apoptotic chromatin compaction remains undefined. To directly compare the extent of histone deacetylation in apoptosis and mitosis, we profiled histone acetylation in HeLa cells undergoing apoptosis or mitotic arrest. We induced apoptosis using Bcl-2 Homology domain 3 (BH3) mimetics^30^, confirming cell death by poly-ADP ribose polymerase (PARP) cleavage, and we synchronised parallel cultures to mitosis, verifying mitotic entry via anti-phospho-Ser/Thr-Pro antibody (MPM-2) staining. Fluorescence-based western blotting showed a marked, global reduction in acetylation across the N-terminal tails of all four core histones in apoptotic cells relative to interphase controls (Fig. 1a–h). The magnitude of this loss was comparable to that observed in mitotic cells, indicating that apoptosis engages a histone deacetylation programme similar to that driving mitotic chromatin condensation.

**Figure 1.**
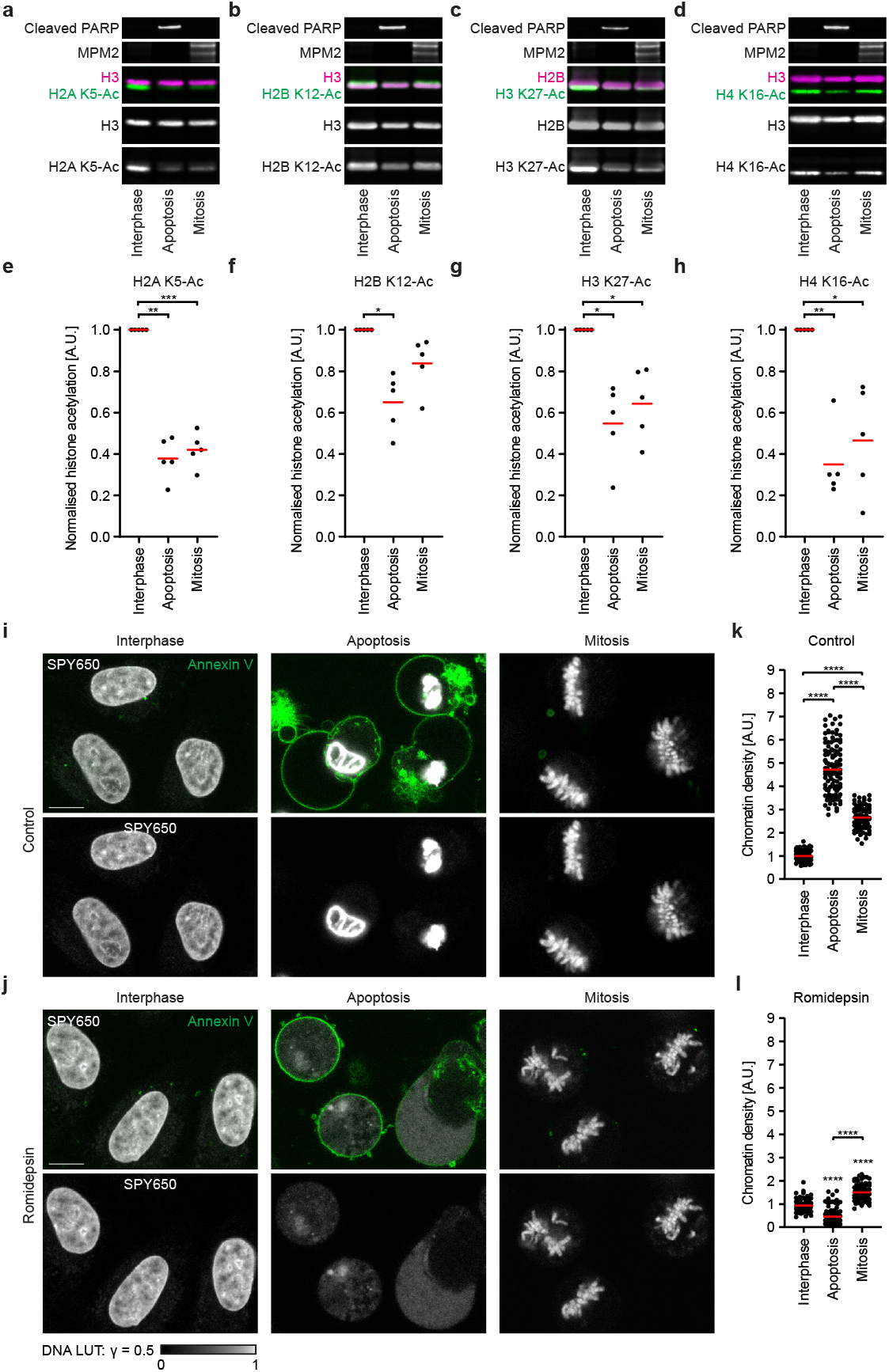
Apoptotic histone deacetylation is required for global chromatin compaction. **a-d**, Fluorescence-based western blot of histone acetylation levels of four core histones during apoptosis and mitosis. Cell lysates were harvested from synchronous interphase, apoptotic and mitotic cultures, and levels of histone acetylation were measured relative to core histones. Cleaved PARP and MPM2 were blotted to detect apoptotic and mitotic states, respectively. **e-h**, Quantification of histone acetylation levels, relative to core histone amounts in the same loading lane. *n*=5 for all conditions. Bars indicate mean; significance was tested by a parametric ratio paired t-test (H2A K5-Ac, *P*=0.0016 (apoptosis) *P*=0.0007 (mitosis); H2B K12-Ac, *P*=0.0122 (apoptosis) *P*=0.0692 (mitosis); H3 K27-Ac, *P*=0.0294 (apoptosis) *P*=0.0218 (mitosis); H4 K16-Ac, *P*=0.0036 (apoptosis) *P*=0.0498 (mitosis). **i, j**, Imaging of chromatin density in the presence and absence of active histone deacetylases in interphase, apoptosis and mitosis. Chromatin was stained with the DNA dye SPY650, and Annexin V-488 was used to detect loss of membrane asymmetry in apoptosis. Scale bar 10 µm. **k, l**, Quantification of chromatin compaction. *n*=131 for interphase, *n*=124 for apoptosis, *n*=84 for mitosis (control); *n*=105 for interphase, *n*=121 for apoptosis, *n*=95 for mitosis (romidepsin). Bars indicate mean; significance was tested by a two-tailed Mann-Whitney test (control apoptosis vs interphase, *P*=<0.0001; control mitosis vs interphase, *P*=<0.0001; control apoptosis vs mitosis, *P*=<0.0001; romidepsin-treated apoptosis vs control apoptosis, *P*=<0.0001; romidepsin-treated mitosis vs control mitosis, *P*=<0.0001; romidepsin-treated apoptosis vs romidepsin-treated mitosis, *P*=<0.0001).

To test whether histone deacetylation is required for chromatin compaction during apoptosis, we applied the class I-specific histone deacetylase (HDAC) inhibitor romidepsin and analysed its effect on chromatin organisation. Cells treated with romidepsin underwent apoptosis normally, as shown by Annexin V staining and PARP cleavage (Fig. 1i, j; Supplementary Fig. 1a-d), but failed to deacetylate histones (Supplementary Fig. 1a-h). In control apoptotic cells, chromatin compacted into a dense mass within the collapsing nucleus, approximately five times as dense as in interphase and twice as dense as in mitotic cells (Fig. 1i, k). Romidepsin-treated cells displayed pronounced chromatin decompaction, both in apoptotic and in mitotic cells, suggesting that active deacetylases are required for compaction in both settings (Fig. 1i-l). Suppressing deacetylation by two independent approaches, either with the broad-spectrum HDAC inhibitor trichostatin A (TSA) or by overexpression of the histone acetyltransferase p300 catalytic domain, elicited a similar phenotype (Supplementary Fig. 2a-d). Together, these findings identify histone deacetylation as a key regulator of chromatin compaction during apoptosis, revealing a conserved regulatory principle shared with mitotic chromosome assembly.

## Chromatin compaction prevents the dispersal of fragmented DNA

The requirement for histone deacetylation during apoptotic chromatin compaction led us to ask how such compaction contributes to the physical containment of fragmented DNA during cell death. Inhibition of histone deacetylation resulted in markedly lower chromatin density in apoptotic cells than in mitotic cells (Fig. 1j, l), despite comparable degrees of histone acetylation (Fig. 1a-h). This suggests that an additional factor controls chromatin organisation when compaction is lost in apoptotic cells. Given that genomic DNA is cleaved during apoptosis^31,32^, we hypothesised that disruption of intact chromosomes caused by this fragmentation might underlie the more pronounced chromatin decompaction in apoptotic cells. To test this, we knocked out the major apoptotic endonuclease, CAD^12–15^, using CRISPR/Cas9 (Supplementary Fig. 3a). Gel electrophoresis confirmed that CAD is essential for apoptotic genome fragmentation in our model (Supplementary Fig. 3b). Confocal imaging showed that CAD knockout did not affect chromatin density in apoptotic cells (Fig. 2a-c), although chromatin typically formed a peripheral ring rather than the dense central mass characteristic of late-stage apoptosis^22^. This is consistent with previous reports that CAD influences chromatin distribution, but not compaction itself^33,34^.

**Figure 2.**
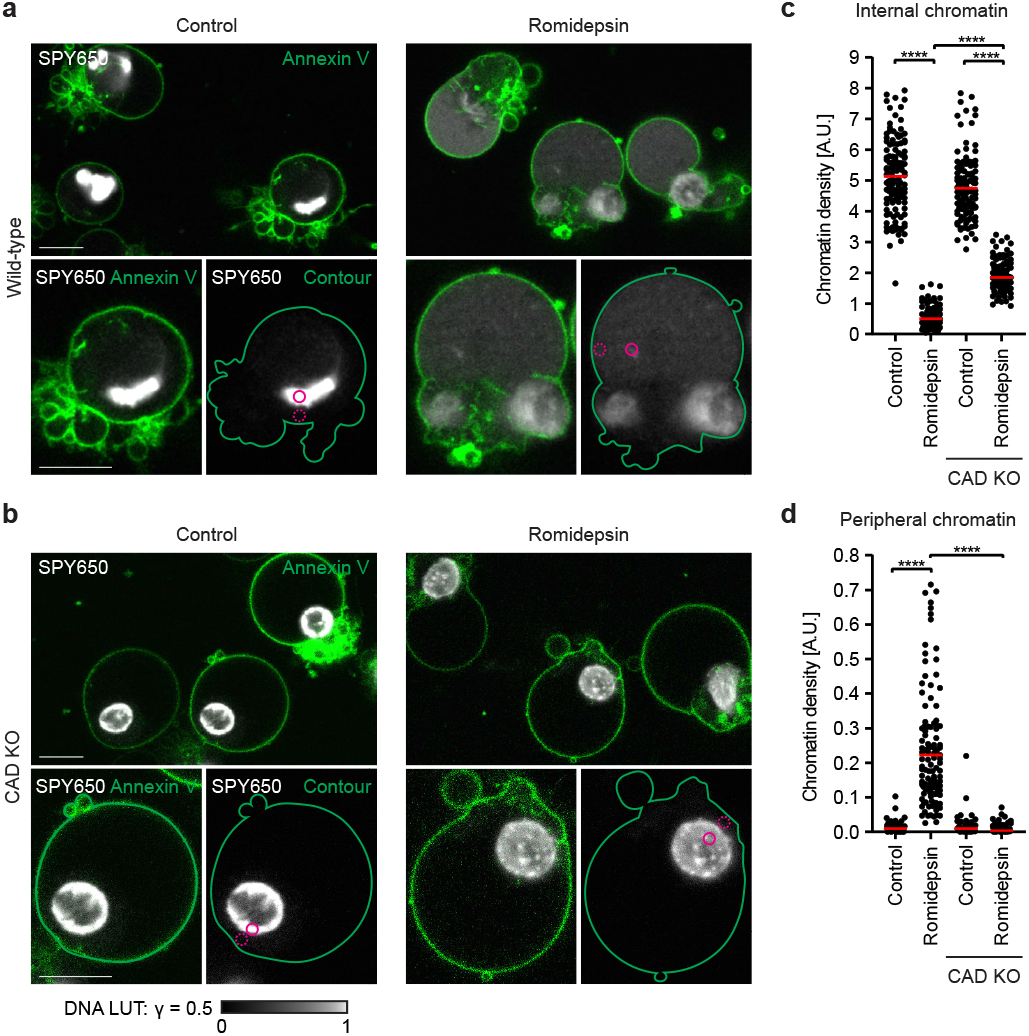
Histone deacetylation prevents dispersion of fragmented apoptotic chromatin. **a, b**, Imaging of chromatin density in the presence and absence of active histone deacetylases in wild-type and CAD KO apoptotic cells. Chromatin was stained with the DNA dye SPY650, and Annexin V-488 was used to detect loss of membrane asymmetry in apoptosis. Contours in the right-hand insets were drawn from the outer contour of Annexin V staining. Scale bar 10 µm, inset 5 µm. Solid circle represents an area of internal chromatin; dotted circle represents an area of peripheral chromatin. **c, d**, Quantification of chromatin compaction at the apoptotic cell interior (**c**) and periphery (**d**). *n*=132 for wild-type apoptosis, *n*=122 for CAD KO apoptosis (control); *n*=116 for wild-type apoptosis, *n*=102 for CAD KO apoptosis (romidepsin). Bars indicate mean; significance was tested by a two-tailed Mann-Whitney test (romidepsin-treated wild-type vs control wild-type, *P*=<0.0001 (internal), *P*=<0.0001 (peripheral); romidepsin-treated CAD KO vs control CAD KO, *P*=<0.0001 (internal); romidepsin-treated CAD KO vs romidepsin-treated wild-type, *P*=<0.0001 (internal), *P*=<0.0001 (peripheral)).

We next asked how genome fragmentation affects chromatin organisation in apoptotic cells that fail to deacetylate histones. In fragmentation-proficient wild-type cells treated with romidepsin, decompacted chromatin spread through the cytoplasm, extending to the plasma membrane (Fig. 2a). In contrast, romidepsin-treated CAD-knockout cells showed only limited chromatin decompaction, with DNA remaining confined to the central cytoplasmic region and never reaching the cell periphery (Fig. 2b). Quantification of DNA intensity across internal and peripheral regions confirmed that chromatin dispersal was strongly reduced in CAD-deficient cells compared with wild-type (Fig. 2a-d). These findings demonstrate that extensive chromatin dispersal in apoptotic cells requires both genome fragmentation and suppression of histone deacetylation. Histone deacetylation-mediated compaction is therefore necessary to prevent chromatin fragments generated by CAD from dispersing throughout the cytosol.

Because the size of chromatin fragments could influence the extent by which DNA disperses through the cytosol, we asked whether chromatin compaction affects the efficiency of CAD-mediated DNA fragmentation. To test this, we compared fragment density in cells with either compacted or decompacted chromatin. Terminal deoxynucleotidyl transferase–mediated dUTP Nick End Labeling (TUNEL) showed similar levels of DNA fragmentation in both control and romidepsin-treated apoptotic cells (Supplementary Fig. 3c-e). Analysis of purified DNA confirmed that fragment size distributions were unaffected by HDAC inhibition in wild-type cells, whereas CAD-knockout cells showed no fragmentation under either condition (Supplementary Fig. 3b). These findings demonstrate that chromatin compaction does not impair CAD activity. Instead, histone deacetylation functions independently of CAD to maintain fragmented DNA sequestered within a single dense chromatin compartment in the dying cell.

## Chromatin compaction safeguards against DNA release into apoptotic vesicles

During apoptosis, plasma membrane blebbing produces ApoEVs that package cellular components for release into the extracellular space, where they disseminate through tissues to influence distal immune responses^5,10,11,20,21^. ApoEVs contain proteins, lipids and RNA, but DNA is strikingly absent from most vesicles^16,17^. The mechanism preventing DNA from entering these vesicles remains unknown. Given that histone deacetylation retains fragmented chromatin in a single dense compartment, we asked how this might affect the DNA content in ApoEVs. To determine how histone acetylation affects the molecular composition of ApoEVs, we profiled their contents by flow cytometry. Annexin V staining was used to discriminate live from apoptotic cells. ApoEVs and apoptotic cell bodies were distinguished by size, and DNA and cytosolic material were detected by labelling with SPY650 and CellTrace Yellow, respectively, and compared to unstained controls (Fig. 3a-f; Supplementary Fig. 4a-c). While nearly all ApoEVs contained cytosolic signal, only 1% showed detectable DNA fluorescence (Fig. 3d-f), consistent with previous studies showing that DNA is selectively depleted from ApoEVs relative to cytosol^16,17^. In contrast, inhibition of histone deacetylation with romidepsin increased the proportion of DNA-positive ApoEVs over tenfold without altering cytosolic content (Fig. 3d-f). These data identify deacetylation-driven chromatin compaction as a key determinant of DNA entry into ApoEVs.

**Figure 3.**
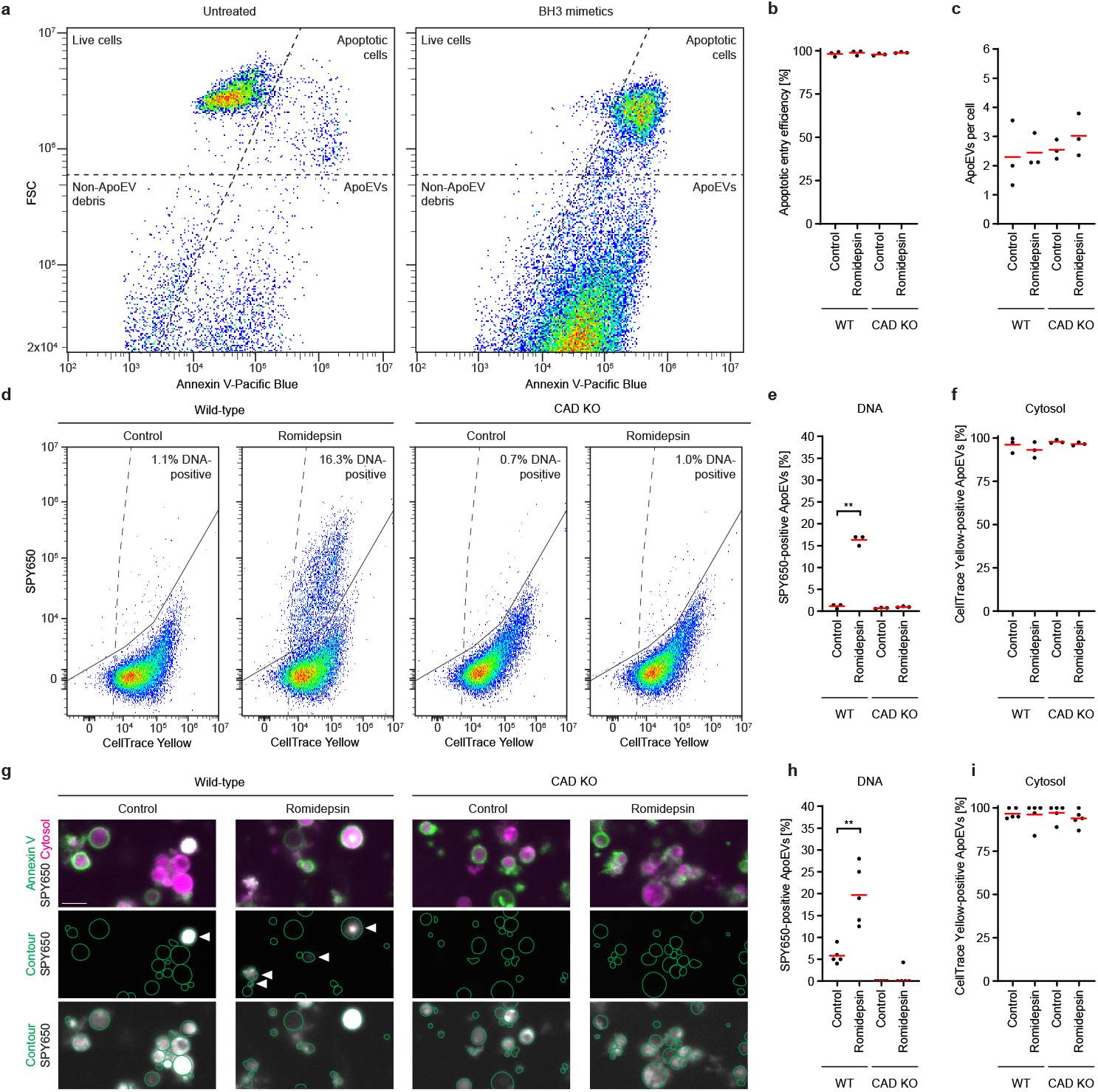
Apoptotic chromatin compaction prevents DNA release into ApoEVs. **a**, Flow cytometry plots of control and BH3 mimetic-treated cells. Apoptotic cells and ApoEVs were detected by Annexin V-488 staining, and ApoEVs were distinguished from cells based on size. **b**, Quantification of apoptotic entry efficiency, the percentage of total cells in apoptosis *n*=3 for all conditions. Bars indicate mean. **c**, Quantification of apoptotic ApoEV production, the number of ApoEVs per apoptotic cell *n*=3 for all conditions. Bars indicate mean. **d**, Flow cytometry plots of DNA and cytosol content in ApoEVs from apoptotic wild-type and CAD KO cells, in the presence and absence of romidepsin. DNA was stained with SPY650 and cytosol with CellTrace Yellow. Solid line represents threshold for positive DNA staining, dotted line represents threshold for positive cytosolic staining. Thresholds were defined by unstained controls. DNA percentages displayed as mean values across three biological replicates. **e, f**, Quantification of ApoEV DNA (**e**) and cytosol (**f**) content, as a percentage of all ApoEVs. *n*=3 for all conditions. Bars indicate mean; significance was tested by a parametric paired t-test (DNA content in romidepsin-treated wild-type vs control wild-type, *P*=0.0031). **g**, Imaging of DNA and cytosol within ApoEVs. ApoEVs were stained with Annexin V-488, DNA with SPY650 and cytosol with CellTrace Yellow. Contours in the lower panels were drawn from the outer contour of Annexin V staining. Scale bar 5 µm. **h, i**, Quantification of imaged ApoEV DNA (**h**) and cytosol (**i**) content, as a percentage of all ApoEVs. *n*=3 for all conditions. Bars indicate mean; significance was tested by a parametric paired t-test (DNA content in romidepsin-treated wild-type vs control wild-type, *P*=0.0039).

To examine how genome fragmentation influences DNA packaging, we analysed ApoEVs from CAD-knockout cells. In contrast to wild-type, romidepsin treatment did not increase the proportion of DNA-containing ApoEVs in CAD-knockout cells (Fig. 3d,e), indicating that hyperacetylated chromatin remains spatially confined when the genome is not fragmented. Notably, neither treatment with romidepsin nor CAD KO induced significant changes in apoptotic entry or ApoEV biogenesis (Fig. 3b,c). These results show that histone deacetylation enables genome fragmentation to proceed in a physically contained manner, preventing DNA fragments from escaping into ApoEVs.

Because ApoEV integrity can be affected by isolation procedures^35^, we verified by imaging that the vesicles we analysed were intact. We separated ApoEVs from larger cell corpses using differential centrifugation^11,36^ and examined the purified fraction by confocal microscopy. Robust Annexin V staining of vesicles smaller than 5 μm in diameter confirmed the high purity of this ApoEV fraction. ApoEVs consistently contained cytosol in all experimental conditions, confirming that these vesicles were structurally intact (Fig. 3g-i; Supplementary Fig. 4d-f). Only romidepsin-treated wild-type cells produced ApoEVs that frequently contained DNA (Fig. 3g-i), confirming that DNA release into ApoEVs requires both genome fragmentation and inhibition of histone deacetylation.

To assess whether this mechanism of DNA sequestration is conserved across cell types, we analysed Jurkat cells, a second well-established human model of apoptosis. Histone acetylation levels declined during apoptosis relative to interphase (Supplementary Fig. 5a-h), and inhibition of histone deacetylation with romidepsin disrupted chromatin compaction and promoted release of DNA fragments into ApoEVs (Supplementary Fig. 5i-l), consistent with observations in HeLa cells. These findings establish histone deacetylation–mediated chromatin compaction as a conserved mechanism that prevents DNA escape from apoptotic cells.

Together, these findings establish histone deacetylation-mediated chromatin compaction as a conserved feature of apoptotic self-disassembly. By coupling genome fragmentation with chromatin compaction, apoptosis dismantles the nuclear genome while preventing the release of fragmented DNA into ApoEVs.

## Synthetic control of chromatin compaction

Our finding that histone deacetylation drives apoptotic chromatin compaction raises the question of how this structural transformation is mediated. Acetylation neutralises the positive charges of lysine residues on histone tails, weakening their electrostatic attraction to negatively charged linker DNA and the acidic patch on neighbouring nucleosomes^37–44^. This physical model predicts that chromatin density could, in principle, be modulated directly by tuning the charge of the nucleosome. However, histone tail modifications also serve as recruitment platforms for a wide range of reader proteins that can alter chromatin architecture through enzymatic or scaffolding activities^45,46^. It therefore remains unclear whether nucleosome charge changes alone are sufficient to promote compaction in apoptotic cells, or whether this process depends on reader-mediated mechanisms.

To test whether nucleosome charge is sufficient to control chromatin density in cells, we developed a synthetic system to modulate nucleosome charge independently of histone modifications. Our approach was designed to mimic the charge effects of histone acetylation or deacetylation while avoiding crosstalk with endogenous chromatin factors. To achieve this, we used green fluorescent protein (GFP) variants carrying defined net surface charges (either +7e or –7e, respectively)^47^ and tethered them with a flexible linker to a nanobody that binds the H2A–H2B dimer interface on the nucleosome^48^. The resulting constructs, Nano-Pos and Nano-Neg, produced charge shifts comparable in magnitude to those caused by changes in histone tail acetylation.

To assess how these synthetic charge tethers affect chromatin organisation, we expressed them in HeLa cells. Nano-Neg expression caused global chromatin decompaction, with dispersion of heterochromatic foci at the nuclear periphery and nucleoli, phenocopying the effects of histone hyperacetylation by HDAC inhibition^49^. In contrast, Nano-Pos induced pronounced chromatin compaction, producing dense speckles enriched at the nuclear periphery and around nucleoli (Fig. 4a, b). Notably, Nano-Pos also restored chromatin compaction in romidepsin-treated cells (Fig. 4a, b), indicating that nucleosome-bound positive charge can induce chromatin compaction regardless of the acetylation status of histone tails. These findings demonstrate that electrostatic modulation of nucleosomes is sufficient to regulate chromatin compaction in intact interphase cells, independently of the effects of histone acetylation.

**Figure 4.**
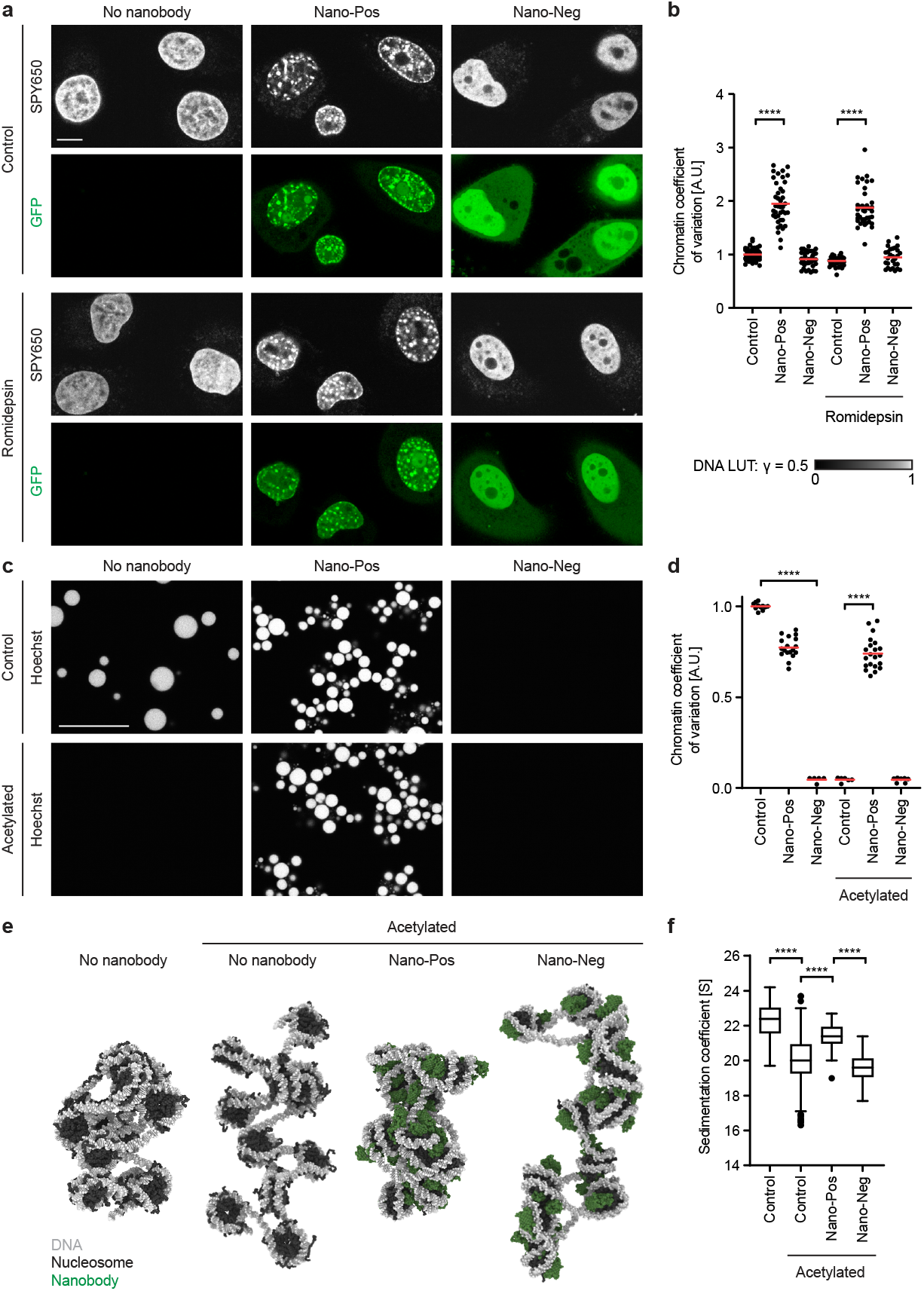
Charge-based synthetic tools to control chromatin compaction. **a**, Imaging of chromatin morphology and nanobody-linked GFP expression in living cells in the presence and absence of romidepsin. Chromatin was stained with the DNA dye SPY650. Scale bar 10 µm. **b**, Quantification of chromatin texture. *n*=69 for control, *n*=41 for Nano-Pos, *n*=36 for Nano-Neg (without romidepsin); *n*=44 for control, *n*=39 for Nano-Pos, *n*=28 for Nano-Neg (romidepsin). Bars indicate mean; significance was tested by a two-tailed Mann-Whitney test (Nano-Pos vs control, *P*=<0.0001 (without romidepsin); *P*=<0.0001 (romidepsin)). **c**, Imaging of naïve and acetylated reconstituted nucleosome arrays in the presence and absence of Nano-Pos and Nano-Neg. Scale bar 5 µm. **d**, Quantification of chromatin phase separation. *n*=11 for control, *n*=18 for Nano-Pos, *n*=5 for Nano-Neg (unmodified); *n*=6 for control, *n*=21 for Nano-Pos, *n*=7 for Nano-Neg (acetylated). Bars indicate mean; significance was tested by a two-tailed Mann-Whitney test (Unmodified Nano-Neg vs unmodified control, *P*=<0.0001; Acetylated Nano-Pos vs acetylated control, *P*=<0.0001). **e**, Simulated folding of a control chromatin fibre and acetylated fibres in the presence and absence of Nano-Pos and Nano-Neg. **f**, Quantification of simulated sedimentation coefficient. *n*=599 for all conditions. Box indicates interquartile range, central bar indicates median, Tukey whiskers shown and outliers included; significance was tested by a two-tailed Mann-Whitney test (acetylated control vs unmodified control, *P*=<0.0001; acetylated Nano-Pos vs acetylated control, *P*=<0.0001; acetylated Nano-Neg vs acetylated Nano-Neg, *P*=<0.0001).

To test whether electrostatic tuning of nucleosomes is sufficient to drive chromatin condensation in the absence of cellular cofactors, we used a biochemical assay based on reconstituted nucleosome arrays. Under physiological salt conditions, unmodified arrays form phase-separated condensates through multivalent nucleosome–nucleosome interactions, whereas histone acetylation disrupts these interactions and keeps arrays dispersed^25,42,44,50^. This minimal system thus provides a platform to assess the physical forces that govern chromatin organisation. We first confirmed that unmodified nucleosome arrays formed condensates under physiological salt, and that acetylation by the histone acetyltransferase p300 prevented condensate formation (Fig. 4c, d), consistent with previous studies^25,42^. We then tested the effect of the synthetic charge tethers. Addition of Nano-Neg to unmodified arrays suppressed condensate formation, mimicking the decompacting effect of acetylation, whereas Nano-Pos restored condensation in acetylated arrays (Fig. 4c, d). These results establish nucleosome electrostatics as a direct physical mechanism for chromatin condensation.

Unlike the flexible histone tails that extend from the nucleosome core, our synthetic effectors present surface charge on a folded protein domain tethered via a flexible linker to the H2A–H2B interface. To investigate how this synthetic charge redistribution influences the balance of attractive and repulsive forces within the chromatin fibre, we performed coarse-grained molecular dynamics (MD) simulations of a 12-nucleosome array^51,52^. Arrays composed of unmodified nucleosomes adopted a compact configuration, whereas acetylation relaxed the fibre into an expanded state (Fig. 4e, f), consistent with previous observations^44^. Simulating the binding of Nano-Pos to acetylated nucleosomes restored fibre compaction to levels comparable to unmodified arrays. By contrast, Nano-Neg failed to reverse the relaxed architecture of acetylated fibres (Fig. 4e, f), consistent with our biochemical and cellular observations. These findings corroborate that when all other chromatin parameters are kept constant, modulation of nucleosome charge alone is sufficient to drive chromatin compaction, even when charge is introduced through an artificial distribution distinct from histone tails.

To gain further mechanistic insight into how Nano-Pos promotes chromatin compaction, we analysed the free-energy landscape of nucleosome-nucleosome interactions across varying distances and orientations. To this end, we calculated the potential of mean force (PMF) as a function of nucleosome separation and relative orientation. In unmodified arrays, nucleosomes engaged in strong attractive interactions across multiple configurations, consistent with efficient fibre condensation. Acetylation markedly reduced this attraction (Supplementary Fig. 6), reflecting disruption of associative electrostatic interactions between nucleosome surfaces. Notably, Nano-Pos restored attraction between acetylated nucleosomes, but primarily when arranged in a side-by-side geometry. In contrast, Nano-Neg did not confer substantial attraction under any configuration. These findings suggest that Nano-Pos promotes compaction through an orthogonal mechanism, re-establishing electrostatic bridging at the nucleosome surface and functionally bypassing the role of histone tails. Together, these results establish that nucleosome electrostatics are sufficient for chromatin compaction in cells and in vitro, independent of histone tail chemistry and binding of reader proteins.

### Electrostatic chromatin compaction sequesters apoptotic DNA

Having established that nucleosome electrostatics are sufficient to compact chromatin in interphase cells, we next asked whether this physical mechanism could also drive changes in chromatin compaction and DNA sequestration during apoptosis. We therefore investigated how the synthetic charge effectors Nano-Pos and Nano-Neg modulate chromatin architecture in dying cells. Expression of Nano-Neg mimicked the effects of HDAC inhibition, producing widespread chromatin decompaction and dispersion throughout the apoptotic cytoplasm. In contrast, Nano-Pos did not disrupt compaction on its own, but when applied to romidepsin-treated cells, it fully restored chromatin compaction (Fig. 5a-c). Thus, synthetic charge tethers can functionally substitute for the endogenous histone deacetylation programme that normally compacts apoptotic chromatin.

**Figure 5.**
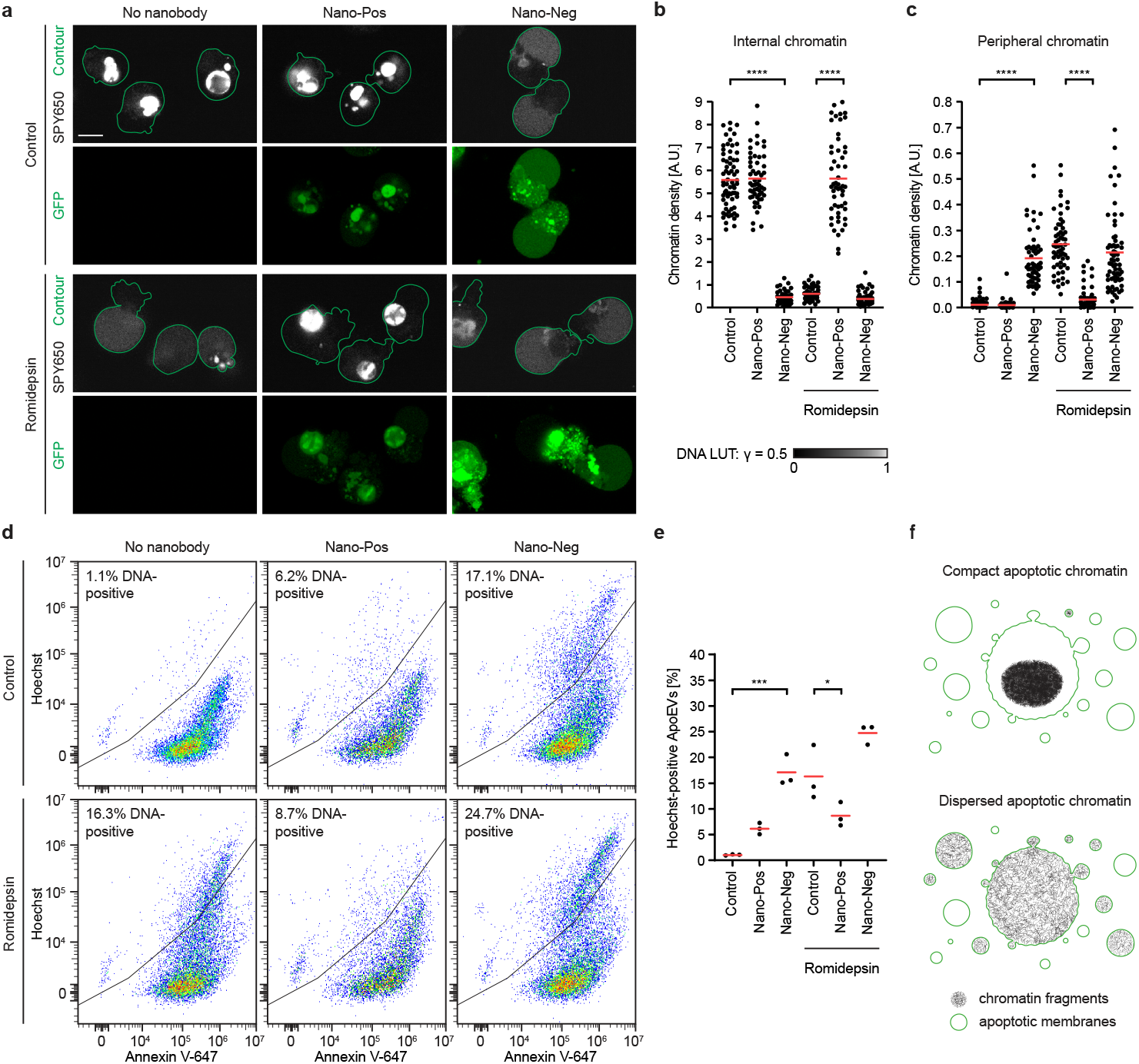
Electrostatics are sufficient to drive chromatin compaction and sequester apoptotic DNA. **a**, Imaging of chromatin density in control and romidepsin-treated apoptotic cells in the presence and absence of Nano-Pos and Nano-Neg. Chromatin was stained with the DNA dye SPY650. Contours were drawn from the outer contour of Annexin V staining. Scale bar 10 µm. **b, c**, Quantification of chromatin compaction at the apoptotic cell interior (**b**) and periphery (**c**). *n*=68 for control, *n*=57 Nano-Pos, *n*=55 Nano-Neg (without romidepsin); *n*=58 for control, *n*=54 Nano-Pos, *n*=68 Nano-Neg (romidepsin). Bars indicate mean; significance was tested by a two-tailed Mann-Whitney test (Untreated Nano-Neg vs untreated control, *P*=<0.0001 (internal), *P*=<0.0001 (peripheral); romidepsin-treated Nano-Pos vs romidepsin-treated control, *P*=<0.0001 (internal), *P*=<0.0001 (peripheral). **d**, Flow cytometry plots of DNA content in ApoEVs from control and romidepsin-treated apoptotic cells, in the presence and absence of Nano-Pos and Nano-Neg. DNA was stained with SPY650 and apoptotic membranes with Annexin V-647. Positive populations were defined by unstained controls. DNA percentages displayed as mean values across three biological replicates. **e**, Quantification of ApoEV DNA content, as a percentage of all ApoEVs. *n*=3 for all conditions. Bars indicate mean; significance was tested by a parametric ratio paired t-test (Untreated Nano-Neg vs untreated control, *P*=0.0005; romidepsin-treated Nano-Pos vs romidepsin-treated control, *P*=0.0233). **f**, Schematic of chromatin compaction and fragmentation during apoptotic cell disassembly.

To investigate how electrostatic tuning by synthetic charge tethers affects the spread of DNA fragments via ApoEVs, we performed flow-cytometric analysis. Nano-Neg increased the proportion of ApoEVs containing DNA, in contrast to Nano-Pos. Nano-Pos also suppressed DNA release under conditions of HDAC inhibition, confirming its ability to restore confinement of fragmented DNA (Fig. 5d–e). These findings demonstrate that direct modulation of nucleosome charge is sufficient to control both chromatin compaction and DNA retention during apoptosis.

## Conclusions

Our findings establish histone deacetylation as a safeguard that sequesters chromatin during apoptosis, preventing the spread of genome remnants via extracellular vesicles (Fig. 5f). This sequestration distinguishes apoptosis from other death pathways: in pyroptosis, chromatin remains diffuse^53^, whereas in NETosis, histone modifications induce extensive decompaction to form extracellular traps that capture pathogens^54,55^. These contrasting outcomes reveal histone acetylation as a universal regulator of how chromatin architecture adapts to distinct cellular fates.

Our results reveal a cooperative mechanism in which genome fragmentation by CAD and chromatin compaction by histone deacetylation together ensure the orderly destruction and confinement of the genome. While CAD-mediated cleavage ensures irreversible destruction of chromosomes, this destruction generates mobile chromatin fragments that could disperse throughout the cytosol and ultimately populate ApoEVs. Deacetylation-driven compaction counterbalances this dispersal by maintaining all chromatin in a single dense compartment, facilitating genome fragmentation without DNA release, even after disassembly of the nuclear envelope. Although the immunological consequences lie beyond the scope of this study, previous work indicates that such sequestration may help preserve the immunological silence of apoptosis by preventing the formation of DNA-rich vesicles, which are known to trigger inflammatory or autoimmune responses in diseases such as systemic lupus erythematosus^20,21^. Modulating chromatin compaction could therefore represent both a physiological apoptotic mechanism for suppressing aberrant DNA-driven inflammation or, conversely, enhance apoptotic immunogenicity in therapeutic contexts.

By reconstituting apoptotic chromatin compaction through a fully synthetic mechanism, we identify nucleosome electrostatics as a minimal physical principle sufficient for genome sequestration. Our programmable charge-based effectors demonstrate that chromatin density can be modulated independently of enzymatic activity or modification-dependent reader proteins, in both interphase and apoptotic cells. Beyond apoptosis, these tools provide a versatile platform to investigate how electrostatic forces interface with histone modification– dependent regulatory codes^45,46^ to shape nuclear organisation and regulate essential processes such as transcription, DNA replication and repair.

By revealing how histone deacetylation couples genome fragmentation to physical chromatin sequestration, this work defines the mechanism that prevents DNA escape during apoptosis. Building on this foundation, future studies should identify the apoptotic signals that trigger histone deacetylation, clarify the contribution of additional apoptotic chromatin-remodelling factors^56–60^, and determine how phagocytes sense or respond to DNA within apoptotic vesicles. By uncovering the interplay between chromatin fragmentation and compaction, and introducing synthetic tools to manipulate nucleosome electrostatics, our study establishes a framework for exploring how charge balance governs genome organisation across cellular states. More broadly, these findings position nucleosome electrostatics as a fundamental physical principle of chromatin organisation, linking biochemical regulation to the material properties that preserve, and ultimately dismantle, the genome.

## Acknowledgements

We thank Daniel Isenberg and David Isenberg for their generous support of the MBL Chromatin Collaborative in honour of their father, Irvin Isenberg. We thank I. Patten for comments on the manuscript. We thank C. Bell, M. Petrovic and M.W.G. Schneider for technical support. This work was further supported by the IMBA / IMP / GMI BioOptics Facility and Protein Technologies Facility at the Vienna BioCenter Core Facilities (VBCF).

## Author contributions

M.F.D.S. and D.W.G. conceived the study. M.F.D.S. performed all cell biological experiments except TUNEL experiments, which were performed by N.S. N.S. performed genome editing to generate CAD KO cell lines. S.W. and M.F.D.S. performed ApoEV experiments. J.H., M.J.M. and J.I.P.L. performed modelling and computer simulations. M.A. and L.C. performed nucleosome array experiments. M.F.D.S. developed image analysis procedures. D.W.G., R.C.-G. and M.K.R. acquired funding and supervised the project. M.F.D.S. and D.W.G. wrote the initial manuscript.

## Materials and methods

### Cell culture

HeLa Kyoto (S. Narumiya, Kyoto University, Japan) and Jurkat E6-1 (ATCC TIB-152) cells were maintained in a 37 °C humidified atmosphere containing 5% CO_2_. HeLa cells were cultured in Dulbecco’s Modified Eagle Medium (DMEM, Gibco) supplemented with 10% (v/v) foetal bovine serum (FBS, Gibco), 1% (v/v) GlutaMAX (Thermo) and 1% (v/v) penicillin–streptomycin (Sigma). Jurkat cells were cultured in Roswell Park Memorial Institute (RPMI) 1640 Medium (Gibco) supplemented with 10% (v/v) foetal bovine serum (FBS, Gibco), 1% (v/v) GlutaMAX (Thermo), 1% (v/v) penicillin–streptomycin (Sigma) and 1 mM Sodium Pyruvate (Sigma). For imaging experiments without cell fixation, HeLa and Jurkat cells were maintained in DMEM, no phenol red (Gibco) and RPMI 1640 Medium, no phenol red (Gibco), respectively. When washing was required, non-adherent Jurkat cells were pelleted by centrifugation at 300 g for 5 min. All cells were regularly tested for the presence of mycoplasma infection, for which no positive results were returned.

### Generation of CAD KO cell lines

Monoclonal CAD KO HeLa and Jurkat cell lines were generated by a Cas9-induced deletion. HeLa cells were seeded in a 24-well plate and transfected the following day with paired gRNAs using Lipofectamine CRISPRMAX Cas9 Transfection reagents (Thermo Scientific), according to manufacturer’s guidelines, resulting in transfection of 1.13 µg Cas9 protein with 0.21 µg respective sgRNAs each per well. For Jurkat cells, an RNP mix consisting of 8.67 µg Cas9 and 1.3 µg of each sgRNA in resuspension buffer was electroporated using Neon Transfection System (Invitrogen) at 1600 V, 10 ms and 3 pulses. Electroporated cells were resuspended in fresh RPMI 1640 medium mixed 1:1 with conditioned medium. HeLa and Jurkat cells were incubated for 48 h before diluting for monoclonal expansion. CAD KO clones were identified by PCR for the absence of an exon region flanked by the gRNA pair. Clonal characterisation included assessment of growth rate and of DNA fragmentation following apoptotic induction.

### Inhibitors and small molecules

To induce apoptosis, cells were treated with a BH3 mimetic cocktail^30^ of 0.5 µM S63845 (Selleck Chemicals, S8383) and 5 µM ABT-737 (Selleck Chemicals, S1002). Apoptotic cells were imaged, fixed or harvested 6 h after induction of apoptosis. To obtain mitotic cells, cells were first synchronised to the G1/S phase boundary by incubating for 16 h in 2 mM Thymidine (Sigma, T1895). Thymidine was removed by washing three times with PBS and cells were allowed to proceed through S phase for 8 h before being arrested at the G2/M boundary by addition of 10 µM RO-3306 (Sigma, SML0569). After 4 h, cells were released into mitosis by washing four times with warm DMEM, while cells were kept on a pre-warmed heat block. After 30 min, mitotic cells were obtained by shake-off and immediately imaged or harvested for western blot. To inhibit HDACs, cells were treated with 50 nM Romidepsin (Selleckchem, S3020) or 500 nM TSA (Sigma, T8552). HADC inhibitors were added 4 h before BH3 mimetic treatment or RO-3306 washout, for apoptotic cells and mitotic cells, respectively.

### Western blotting

Cells were seeded in 6-well plates at a density of 500 k cells (HeLa) or 1.2 million cells (Jurkat) per well of 6-well plates. Mitotic synchronisation and apoptotic induction were performed as described above, and cells were harvested a day after seeding. For harvesting, interphase HeLa cells were trypsinised and resuspended in warm DMEM. All cells were centrifuged for 4 min at 125 g, before being resuspended in 1 ml ice cold PBS. Cells were kept on ice and counted, and a volume corresponding to 700 k HeLa cells or 2 million Jurkat cells was centrifuged for 4 min at 125 g, at 4°C. The pellet was resuspended in 100 µL 6x Laemmli buffer (375 mM Tris pH 6.8 (Sigma); 12% SDS w/w (Serva); 60% Glycerol w/w (Sigma); 0.06% Bromophenol blue (Merck)), supplemented with 60 mM DTT (Sigma) and 1 U Benzonase (Merck). Samples were incubated at room temperature for 5 min, then boiled at 95°C for 10 min. 3.5 µL of each sample was loaded in 10-well 1.0 mm 4-12% NuPage Bis-TRIS gels (Thermo), alongside 3.5 µL Chameleon™ Duo Pre-stained Protein Ladder (Li-COR, 928-60000). Gels were run in MES SDS Running Buffer (Thermo) at 120 V for 10 min, then at 200 V for 20 min. Gels were removed and equilibrated in Transfer Buffer (20% ethanol w/w (Sigma); 10% 10X Transfer Buffer (Thermo)) for 10 min. Proteins were transferred onto 0.2 µm Amersham™ Protran nitrocellulose membranes (Cytiva, 10600001) using the Mini Trans-Blot® Cell system (BioRad). After transfer, membranes were blocked 1 h at room temperature in 5% filtered Bovine serum albumin (Sigma) in 0.05% T-BST (50 mM Tris-HCl pH, 7.5 (Sigma), 150 mM NaCl (Sigma), 0.05% Tween 20 (Sigma). Membranes were incubated in primary antibody in 5% BSA / 0.05% TBST on a rocker overnight at 4°C, before three brief rinses in 0.05% TBST and three 5 min washes, also in 0.05% TBST. Membranes were then incubated in secondary antibody in 5% BSA / 0.05% TBST for 1 hr at room temperature, before three brief rinses in 0.05% TBST and three 10 min washes, also in 0.05% TBST. Stained membranes were imaged using a ChemiDoc MP Imaging System (BioRad). Histone acetylation was quantified in Fiji, with each acetylated epitope of interest normalised to fluorescence levels of another core histone, detected on the same membrane by detection with IRDye 680RD or IRDye 800CW-conjugated secondary antibodies (Li-COR).

### Cell and ApoEV microscopy

Before imaging, HeLa cells and Jurkat cells and ApoEVs were seeded into Lab-Tek 8-well Chambered Coverglass slides (Thermo; 155409) at densities of 25 k cells (HeLa) or 60 k cells (Jurkat) per well. The next day, cells were treated with romidepsin and BH3 mimetics as described above. 3 h before imaging, cells were stained by addition of 1:50 Annexin V-Alexa Fluor™ 488 (Thermo, A13201) or 1:50 Annexin V-Pacific Blue™ (Thermo, A35122) to detect apoptotic membranes; and 1:1,000 SPY650-DNA (Spirochrome, SC501). For nanobody-expressing cells, samples were stained with 1.62 µM Hoechst 33342 (Invitrogen, H1399) to avoid crosstalk with fluorescent proteins. All stains were added in imaging media in a 10X staining mix to prevent disruption of apoptotic and mitotic cells. Cells were imaged using a customised Zeiss LSM780 microscope, using a X63 1.4 numerical aperture oil-immersion DIC plan-apochromat objective (Zeiss) and ZEN Black 2011 software. For imaging experiments without fixation, cells and ApoEVs were maintained at 37°C and 5% CO_2_ during image acquisition using a customised incubation chamber (EMBL). A custom aluminium mounting block (IMP/IMBA workshop and BioOptics) was used to mount Lab-Tek imaging chambers.

### Gel electrophoresis of apoptotic DNA fragments

To isolate fragmented DNA after apoptotic induction, 400 k HeLa cells or 800 k Jurkat cells were seeded into 6-well and 12-well plates, respectively. The following day, respective wells were treated with romidepsin and BH3 mimetics as described above. 700 k cells (HeLa) and 500 k cells (Jurkat) were harvested by trypsinisation, pelleted and then washed once with ice-cold 1X PBS prior to counting. After counting, cells were pelleted at 300 g at 4°C. Pellets were resuspended in DMSO and vortexed briefly, then TE-2% SDS buffer was added and vortexed briefly. Samples were centrifuged at 16,000 g for 10 min at 4°C. The supernatant was mixed with DNA binding buffer and passed through a silica column. DNA was eluted in TE buffer and incubated at 65°C for 90 min. Samples were electrophoresed in 2% agarose gel at 120V for 120 min with GelRed (Biotium) DNA stain.

### Immunofluorescence and TUNEL assay for microscopy with cell fixation

Lab-Tek 8-well Chambered slides were treated with poly-L-lysine (Sigma) before HeLa cells were seeded at a density of 25 k cells per well. After induction of apoptosis, cells were fixed with 4% paraformaldehyde in 1X PBS, quenched with 10 mM TRIS-HCl, and permeabilized with 0.2% TritonX-100. For immunofluorescence, wells were blocked with 5% BSA in PBS for 30 min with rocking before primary antibody Cleaved PARP (Asp214) (D64E10) XP® Rabbit (Cell Signaling Technologies; #5625S; Lot 18) was added at 1:400 dilution for 2 h with rocking. After washing 3x with 1X PBS, Hoechst 33342 was added at 1:10,000 dilution and secondary antibody Donkey anti-rabbit IgG (H+L) Highly Cross-Adsorbed Alexa Fluor™ 594 (Thermo; #A-21207, lot 1744751) was added at 1:1,000 dilution for 1 hour with rocking. Cells were washed 3x with 1X PBS. Click-iT™ Plus TUNEL Assay Kit for In Situ Apoptosis Detection Alexa Fluor™ 488 (Thermo; #C106179) was used to detect DNA fragment ends. The control well was washed once with ddH_2_O and then treated with 2U DNase I for 30 min at room temperature. All wells were washed once with ddH_2_O, then TdT Reaction Buffer was added and incubated at 37°C for 10 min. TdT Reaction mixture was added to each well and incubated at 37°C for 60 min then washed with 5% BSA in PBS twice for 7 min and once with 1X PBS. Click-iT Plus TUNEL reaction cocktail was added to all wells and incubated at 37°C for 30 min then washed with 5% BSA in PBS twice for 7 min and once with 1X PBS. Cells were then imaged by confocal microscopy.

### Image analysis

Chromatin density and TUNEL density in live and apoptotic cells was recorded in Fiji by measuring three random regions of interest (ROIs) each with 600 nm diameter, within a central z-section of the cell. Values for each cell were taken as the mean intensity of SPY650 or TUNEL within each ROI, and corrected for background fluorescence integrity. Background-subtracted values were normalised relative to control interphase cells. For peripheral chromatin measurements, ROIs were restricted to a region within and 1 µm of the apoptotic cell membrane. For coefficient of variation measurements, standard deviation and mean intensity was recorded in Fiji by measuring three random ROIs each with 6.4 µm diameter, within a central z-section of the cell. Coefficient of variation was calculated by dividing standard deviation by mean intensity for each ROI. Values for each cell were taken as the mean coefficient of variation of each ROI, and normalised relative to control interphase cells. For ApoEV imaging experiments, mean intensity was recorded in Fiji by measuring one ROI with 1.4 µm diameter, within a central z-section of the ApoEV. SPY650 and CellTrace Yellow mean intensity for each ApoEV was normalised to unstained controls, and ApoEVs were considered to be stained positive if normalised mean intensity values were >1.5x unstained signal. For nucleosome array imaging experiments, Hoechst standard deviation and mean intensity was recorded in Fiji by measuring whole fields-of-view. Coefficient of variation was calculated by dividing standard deviation by mean intensity for each ROI. Values for each experimental repeat were taken as the mean coefficient of variation of each field-of-view, and normalised relative to the control no nanobody condition. To facilitate the comparison of chromatin morphologies and TUNEL signal between control and romidepsin-treated cells, images were displayed on a look-up table (LUT) with an adjusted gamma (*γ*) of 0.5. This adjustment allows for the clear visualisation of DNA distributions across a dynamic range spanning an order of magnitude in brightness.

### Preparation of samples for ApoEV flow cytometry analysis

Cells were seeded in 96-well plates at a density of 20 k cells (HeLa) or 100 k cells (Jurkat) per well. The next day, cells were treated with romidepsin and BH3 mimetics as described above. Before BH3 mimetic treatment, cells were loaded with 5 µM CellTrace Yellow. Loading dye was removed by washing three times with fresh media. 9 h after apoptotic induction, cells and ApoEVs were pelleted by centrifugation at 20,000 g for 20 min. For live HeLa controls, cells were washed with PBS and trypsinised before harvesting. During centrifugation, staining buffer was prepared in a 5-fold dilution of 5X Annexin V binding buffer (Thermo) in dH_2_O. Samples were stained by addition of 1:50 Annexin V-Alexa Fluor™ 488 (Thermo, A13201) or 1:50 Annexin V-Pacific Blue™ (Thermo, A35122) to detect apoptotic membranes; and 1:1,000 SPY650-DNA (Spirochrome, SC501). Unstained and single-stained samples were prepared for compensation, and full-minus-one (FMO) samples for assessment of background staining. Samples were resuspended in 65 µL of each relevant staining buffer, and transferred to a FACS-compatible V-bottomed 96-well plate (Nunc).

### Flow cytometry

Flow cytometry analysis was performed on an Agilent NovoCyte Penteon five laser flow cytometer, with a core diameter of 7.7 µm and flow rate of 14 µL/min. Mixing was performed for 10 s at 1,500 rpm before acquisition of each sample.

### Flow cytometry compensation and analysis

Flow cytometry data was analysed using FlowJo v10.10.0. For each biological replicate, compensation was independently applied using single-stained and unstained samples. Empty samples were used to set an FSC-H threshold of 20,000 for events above electrical interference. Apoptotic cells and ApoEVs were assigned based on elevated Annexin V staining, relative to live controls. ApoEVs were distinguished from apoptotic cells based on FSC-H intensity. Positive gates for SPY650 and CellTrace Yellow were defined based on FMO-stained control samples. For nanobody experiments, ApoEVs from nanobody-expressing cells were detected y gating for ApoEVs stained positive for E2-Crimson. All plots displayed represent concatenates across biological repeats, equalised to display consistent numbers of total events across contrasting experimental conditions.

### Isolation of ApoEVs for imaging

Cells were seeded in 6 cm dishes at a density of 1 million cells per well. The next day, cells were treated with romidepsin and BH3 mimetics and loaded with CellTrace Yellow as described above. 9 h after apoptotic induction, ApoEVs were isolated by differential centrifugation by an adapted protocol to that previously described^36^. Cells were pelleted out by two rounds of centrifugation: once at 125 g for 5 min (gentle) to remove the majority of cells while preventing excessive cell rupture; before the supernatant was centrifuged a second time at 300 g for 5 min (moderate) to remove residual cells. ApoEVs were pelleted from the resulting supernatant by centrifugation at 2,000 g for 20 min (hard), before ApoEVs were resuspended in 200 µL DMEM, no phenol red (Gibco) containing 1:50 Annexin V-Alexa Fluor™ 488 (Thermo, A13201) or 1:50 Annexin V-Pacific Blue™ (Thermo, A35122) to detect apoptotic membranes; and 1:1,000 SPY650-DNA (Spirochrome, SC501). Samples were transferred to a Lab-Tek 8-well Chambered slide and ApoEVs were left to settle for 20 min before imaging.

### Transfection of plasmids and FACS enrichment

For expression by transient transfection of nanobody or p300 constructs in HeLa cells, 10 cm plates were transfected with a transfection mixture containing 1.5 mL Opti-MEM™ I Reduced Serum Medium (Gibco), 45 µL X-tremeGENE™ 9 DNA Transfection Reagent (Roche), according to manufacturer’s guidelines. For flow cytometry experiments, Nano-Pos and Nano-Neg constructs were cloned into polycistronic vectors containing a P2A self-cleaving polypeptide sequence, allowing stoichiometric expression of the nanobody of interest and an E2-Crimson fluorescent reporter protein. For imaging of nanobody- or p300-expressing cells, FACS was performed 28 h after transfection, gating for fluorescently positive populations relative to a mock transfected control. Positive cells were then seeded into Lab-Tek 8-well Chambered slides prior to romidepsin treatment, apoptotic induction, staining and imaging.

### *In silico* chromatin modelling

Chromatin was simulated using our chemically specific coarse-grained model^51^. In this model, each amino acid is represented by a single bead that retains the amino-acid– specific hydrophobicity, charge, and size of its atomistic counterpart. The secondary structure of the histone globular domains is preserved using a Gaussian elastic network model, while the histone tails are treated as fully flexible polymers. DNA is represented using a modified Rigid Base Pair model, in which each base pair is treated as an ellipsoid decorated with two phosphate charges. Sequence-dependent mechanical properties of DNA are captured by parameterising the six intra–base-pair helical displacements from large-scale atomistic simulations.

The Nano-Pos and Nano-Neg nanobodies were generated using AlphaFold^61^. The nanobodies were manually positioned onto the histone core using VMD to achieve the desired binding geometry, and the resulting complexes were stabilised by introducing covalent bonds when coarse-graining the structure between selected residues at the binding interface.

MD simulations were performed in LAMMPS (version 3 March 2020), using our custom code. Simulations of the chromatin fibres were performed using Debye-Hückel replica exchange, with 16 replicas for each simulation condition^51^. The simulations were run at a constant temperature of 300 K using the Langevin thermostat. The Debye-Hückel screening parameter κ was varied across the replicas, from 8 Å-1 to 15 Å-1, with spacings optimised to ensure an exchange rate near 0.3. All systems were simulated for a total of 1 µs, with a timestep of 10 fs. The initial 200 ns were discarded from the analysis to allow for system equilibration.

#### Sedimentation Coefficient

The sedimentation coefficient *s* was computed using a custom Python implementation of the HullRad method^62^. The coefficient is given by

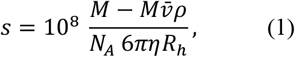

where *M*is the total molar mass, 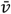 is the partial specific volume, *ρ* is the solvent density, *η* is the solvent viscosity, *R*_*h*_ is the hydrodynamic radius, and *N*_*A*_ is Avogadro’s number. The prefactor converts the sedimentation coefficient to Svedberg units (S).

The total molar mass was computed as

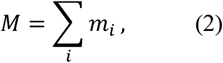

and the partial specific volume was calculated as the mass-weighted average

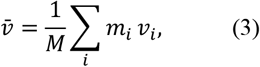

where *m*_*i*_ and *v*_*i*_ are the mass and specific volume of residue (or bead) *i*, respectively.

The hydrodynamic radius *R*_*h*_was estimated from the convex hull of the chromatin structure. To define the molecular envelope, only the coordinates of the DNA beads were used to compute the convex hull. The resulting hull was expanded by a 2.8 Å hydration shell, following the standard HullRad treatment. The hydrodynamic radius was then taken as the radius of a sphere with the same volume as the hydrated convex hull.

#### PMF calculations

Due to the disk-like geometry of the histone octamer, pairwise interactions were decomposed into three orientation-dependent configurations: face–face, face–side, and side–side. For each configuration, a separate PMF was computed using umbrella sampling.

An initial single-nucleosome structure was first equilibrated and then duplicated to form a two-nucleosome system. The two nucleosomes were initially placed with a center-to-center separation of ∼200-240 Å. They were then rotated into the desired relative orientation and maintained in that configuration using the COLVARS package in LAMMPS^63^. A harmonic restraint with a force constant of 1 kcal·mol^−1^·deg^−2^, centered at 0°, was applied to fix the relative orientation of the histone cores.

Initial configurations for the umbrella sampling windows were generated using steered molecular dynamics (SMD), in which *R* was reduced to 70 Å over 1 million timesteps, using a harmonic pulling potential with a force constant of 0.01 kcal·mol^−1^·Å^−2^. The resulting range of *R* was divided into equally spaced windows with separations of 5 Å. Each window was simulated for 10 million timesteps under a fixed harmonic bias centred at the target value of *R*, with a force constant of 0.05 kcal·mol^−1^·Å^−2^. Orientational restraints were maintained throughout all umbrella sampling simulations using the same parameters as in the setup stage.

The PMFs for each interaction geometry were reconstructed from the biased trajectories using the weighted histogram analysis method (WHAM)^64^.

### Construction of the polycistronic histone expression plasmid

A polycistronic plasmid encoding the four human core histones for expression in E. coli (H2A, H2B, H3.1, and H4) was kindly provided by Carrie Bernecky (ISTA, Klosterneuburg). The vector is based on the pET-29a backbone. The plasmid was further modified due to low expression level of H3.1 and H4 histones. For enhanced expression of both histones, additional T7 promoter elements were introduced immediately upstream of the H3.1 and H4 open reading frames, and coding sequences were codon-optimised for E. coli using VectorBuilder. In the final design, H2A carries an N-terminal S-tag–6×His tag followed by a TEV site, and H3.1 carries an N-terminal 6×His tag followed by a TEV site. Additionally, to enable tag removal without introducing non-native residues, the TEV protease recognition sites in the H2A and H3.1 fusion constructs were changed to ENLYFQM such that TEV cleavage leaves an N-terminal methionine as the only residual residue (i.e., no additional ectopic amino acids). The modified and optimised DNA elements were synthesised by GenScript as a gene fragment and replaced the original sequences by restriction digestion and Gibson assembly.

### Expression and Purification of human histones

#### Expression

Expression and purification were performed as previously described^65^ with a few modifications. The optimised plasmid carrying all four histones was transformed into E. coli BL21 DE3 pLysS strain (MolBio Service, VBC) using the heat shock method and plated on Kanamycin LB agar plates (Media Kitchen, VBC) and left to incubate at 37°C overnight. On the following day, 15-20 colonies were picked from the plate and resuspended in 600 µL LB medium. 100 µL of the resuspended cells were added to 1 L 2xTY in a 5 L flask supplemented with Chloramphenicol (25 µg/mL final concentration) and Kanamycin (50 µg/mL) and left on the bench overnight without shaking to minimise the number of generations. On the following day, the flasks were shifted to 37°C and shaking at 180 rpm until they reached OD of 0.4. Protein expression was induced via the addition of 0.4 mM IPTG (Isopropyl β-D-1-thiogalactopyranoside) and left to express at 37°C overnight. On the following day, cells were harvested via centrifugation at 6,000 g for 20 min and then resuspended in cold 1x PBS and transferred to 50 mL falcon tubes. Cells were then centrifuged for 20 min at 3,000 g and the supernatant was discarded and dried pellets were flash frozen in liquid nitrogen and stored at -70°C until purification.

#### Purification

Purification was carried out under non-denaturing conditions. Cell pellets were thawed on ice for 1 h and then resuspended in cold lysis buffer (5 mL/g pellet) using a Polytron PT 2500 homogeniser (lysis buffer: 20 mM Tris-HCl pH 8.0, 2.0 M NaCl, 30 mM imidazole, 1 mM DTT, 1 mM PMSF, lysozyme to a final concentration of 0.5 mg/mL, Roche EDTA-free protease inhibitor cocktail, and 0.1% Triton X-100). The suspension was sonicated using a Branson sonifier (40% amplitude, 2 s ON/3 s OFF, 2 min), incubated on ice for 10 min, and sonicated a second time under the same conditions. The lysate was centrifuged at 40,000 g for 1 h at 4°C, and the supernatant was filtered through a 0.45 µm PVDF membrane. The cleared lysate was diluted 1:10 in binding buffer (20 mM Tris-HCl pH 8.0, 2.0 M NaCl, 30 mM imidazole, 1 mM DTT) to reduce DNA binding and thus contamination before loading onto a HisTrap column.

The diluted lysate was loaded onto a pre-equilibrated Cytiva HisTrap FF 5 mL column at 2 mL/min using an ÄKTA Pure system, followed by washing with 10 column volumes (CV) of binding buffer. Histones were eluted using a stepwise gradient of elution buffer (20 mM Tris-HCl pH 8.0, 2.0 M NaCl, 500 mM imidazole, 1 mM DTT) as follows: 7% elution buffer (5 CV), 20% (5 CV), 40% (5 CV), 60% (5 CV), and 100% (5 CV). Histones typically eluted at 20-40% elution buffer. Histone-containing fractions were pooled, and the total amount was estimated by absorbance at 280 nm. Super-TEV protease (Protein Technologies, VBC) was added at a ratio of 2 mg histone to 1 mg protease. A high protease amount was used because the high salt concentration reduces cleavage efficiency. The sample was dialysed overnight against dialysis buffer (20 mM Tris-HCl pH 8.0, 2.0 M NaCl, 5 mM DTT).

The next day, cleavage was confirmed to be complete by SDS-PAGE. The dialysed sample was loaded onto a Cytiva HisTrap FF 5 mL column (pre-equilibrated in dialysis buffer) at 2 mL/min. The column was washed with 5 CV dialysis buffer, then with 7 CV of 10% elution buffer, 7 CV of 15% elution buffer, and 7 CV of 100% elution buffer. Most histones eluted in the 10% wash step; therefore, the flowthrough and the other fractions were discarded.

Pooled fractions were concentrated to 500 µL using Amicon Ultra 10 kDa MWCO concentrators and loaded onto a Cytiva Superdex 200 pg 16/60 HiLoad size exclusion column pre-equilibrated in SEC buffer (20 mM Tris-HCl pH 8.0, 2.0 M NaCl, 1 mM DTT, 1 mM EDTA) using a 1 mL loop. SEC peaks were analysed by 20% SDS-PAGE, and the peak showing histones in an equimolar ratio was pooled. The sample was concentrated to 5 mg/mL (measured by UV absorbance at 280 nm using a quartz cuvette), aliquoted (50 µL) in Eppendorf Protein LoBind tubes, flash frozen in liquid nitrogen, and stored at −80°C. The purity is >95%.

This procedure yielded ∼2 mg of purified histone octamer per litre of bacterial culture.

### Preparation of 12×601 DNA array for *in vitro* chromatin assembly

Histone octamer assembly on DNA was first tested using a 191 bp double-stranded DNA fragment containing the 601-nucleosome positioning sequence. The DNA was PCR-amplified and gel-purified. Nucleosomes were assembled at room temperature using the dilution method (microscale reconstitution) as described previously^66^. Histone octamer was titrated against 1 µg DNA at histone:DNA molar ratios of 0.8:1, 0.9:1, 1.0:1, 1.1:1, 1.2:1, and 1.3:1.

Assembly reactions were analysed by EMSA on 6% Invitrogen DNA retardation gels running at 100V for 90 min in 0.5xTBE in the cold room. For each reaction, 8 µL of sample was mixed with 2 µL of 5× Hi-Density TBE sample buffer prior to loading. The fraction of assembled nucleosomes was quantified by calculating the ratio of shifted DNA to free DNA. Based on this analysis, a 1.1:1 octamer:DNA molar ratio was used for subsequent assemblies.

Plasmid DNA was prepared from a 3 L E. coli culture using the Invitrogen Gigaprep kit according to the manufacturer’s instructions, yielding ∼15 mg plasmid. For release of the 12×601 array insert, 1 mg plasmid was digested with BbsI-HF and EcoRV-HF (10 µL of each enzyme) at 37°C overnight. Digestion produced a ∼2.1 kb fragment containing the 12×601 repeats with a CCCC overhang, together with multiple fragments <400 bp. The small fragments were retained as carrier DNA to reduce over-assembly of nucleosomes. The digest was purified by two rounds of phenol/chloroform/isoamyl alcohol extraction followed by ethanol precipitation and resuspended in 1× TE buffer to a final concentration of ∼4– 5 µg/µL. DNA concentration was determined by NanoDrop using a serial dilution series and back-calculating the stock concentration.

Nucleosome array assembly was performed as described previously^42^. Briefly, 500 µg of the digested plasmid DNA was adjusted to 2.0 M NaCl by mixing with cold 4 M assembly buffer (10 mM Tris-HCl pH 8.0, 1 mM EDTA, 4.0 M NaCl, 2 mM DTT). The DNA was then diluted with cold 2.0 M assembly buffer (10 mM Tris-HCl pH 8.0, 1 mM EDTA, 2.0 M NaCl, 1 mM DTT) to a final concentration of 601 DNA of 5 µM. Histone octamer was added at an octamer:601 molar ratio of 1.1:1. The reaction was then transferred to a dialysis tubing Spectra/Por 7 pretreated and put in a beaker containing 2L of high salt buffer (10 mM Tris-HCl pH 8.0, 1 mM EDTA, 2 M KCl, 1 mM DTT).

Salt dialysis was carried out at 4°C using a peristaltic pump (0.8 mL/min) to exchange into low-salt buffer (10 mM Tris-HCl pH 8.0, 1 mM EDTA, 0.2 M KCl, 1 mM DTT). After completion of the exchange (∼42 h), the reaction was transferred to no-salt buffer (10 mM Tris-HCl pH 8.0, 1 mM EDTA, 1 mM DTT) and dialyzed overnight.

The assembled arrays were purified by sucrose gradient centrifugation. A 12 mL 15–40% (w/w) sucrose gradient was prepared in the same buffer using a BioComp Gradient Station, and samples were centrifuged in a Sorvall Discovery 90SE with a TH-641 swinging-bucket rotor at 30,000 rpm (154,000 g) for 16 h at 4°C using a slow acceleration and no brake. Gradients were fractionated manually into 200 µL fractions. Protein-containing fractions were identified by mixing 10 µL of each fraction with 2 µL Bio-Rad Protein Assay Dye Reagent Concentrate. DNA content was assessed by treating 5 µL of each fraction with 1 µL Qiagen Proteinase K, incubating at 55°C for 30 min, and analysing samples on a 1% agarose gel stained with SYBR Safe. High-purity fractions were pooled and dialysed against 2 L of no-salt buffer overnight, ensuring sufficient headspace in the dialysis tubing as the sample volume expands during this step. The next day, the buffer was replaced with fresh 2 L no-salt buffer and dialysis was continued overnight. The nucleosome array was typically found at 30% sucrose.

Finally, the sample was concentrated to approximately half of the original volume using a Pierce 10 kDa MWCO concentrator. To determine array concentration, 2 µL of the sample was treated with Qiagen Proteinase K, purified using a DNA purification kit (MolBio Service, VBC), and DNA concentration was used to calculate nucleosome array (and nucleosome) concentration. Quality control of the arrays was then performed via MNase digestion and EMSA after cleaving the arrays with EcoRI-HF, which cleaves all linkers.

### Labelling Array DNA

To fluorescently label the nucleosome arrays for subsequent imaging we leveraged the CCCC overhang created by the restriction digestion. Two short complementary oligos with one carrying a phosphorylated 5’ GGGG overhang and the other carrying a 5’ Atto 647N (Microsynth) were annealed by mixing to a final concentration of 10 µM of each in 0.5xPBS and then heating to 95°C for 2 min, reducing temperature to 80°C, and then gradually decreasing the temperature further to 20°C over the course of 6 h using a BioRad Thermocycler.

For the ligation reaction, Nucleosome arrays were mixed with the annealed oligos at a ratio of arrays to oligos 1:5 in a final reaction buffer of (50 mM Tris-HCl pH 7.6, 2 mM MgCl_2_, 1 mM ATP, 2 mM DTT.) Then, T7 ligase (NEB) was added, and the reaction was incubated overnight at 16°C.

On the following day, the arrays were purified from the excess oligos and T7 ligase using Amicon mini 0.5 mL concentrators with 100kDa MWCO via three rounds of concentration and dilution in no-salt buffer. Samples were then concentrated to 5 µM of nucleosomes. This procedure removes >99% excess oligos and >70% T7 ligase. No observable loss of chromatin was detected and quality control of ligated arrays via MNase digestion and EMSA revealed that they are indistinguishable from untreated samples.

### Recombinant protein expression and purification of Nano-Pos and Nano-Neg

His-tagged expression constructs were transformed into *E. coli* strain Bl21(DE3), and the pre-culture in LB + kanamycin a (K4000, Sigma-Aldrich) was grown overnight at 37°C. The next day, this culture was diluted 1:250 and supplemented with 1.5% w/v lactose (Milvolis, DM), before incubation with shaking at 220 rpm at 25°C for 24 h. Cell pellets were resuspended in 150 mL of IMAC A buffer (50 mM Hepes (pH 7.5) (A1069, PanReac/AppiChem), 1000 mM NaCl (1,06404m Supelco), 20 mM Imidazole (38994, Roth), 1 mM TCEP (HN95.2, Roth)) supplemented with Benzonase (2 μl per 1 mL of buffer, prepared in-house), 0.15% v/v Igepal CA630 (56741, Supelco) and three tablets of protease inhibitors (Roche, EDTA free), and then lysed by sonication (19 mm probe, 5 min: Pulse on 1s, pulse off 2s, 60% Amplitude). The soluble fraction was isolated by centrifugation at 20,000 g, 30 min at 4°C, then loaded at a flow rate of 2 mL/min onto a 5 mL HisTrap FF column (17525501, Cytiva), previously equilibrated with IMAC A buffer, and the column was finally washed with 120 mL of the same buffer. The column was developed with 20 mL step gradients of 6%, 16%, 21%, 46%, and 100% IMAC B buffer (50 mM Hepes (pH 7.5) (A1069, PanReac/AppiChem), 500 mM NaCl (1,06404m Supelco), 500 mM Imidazole (38994, Roth), 1 mM TCEP (HN95.2, Roth)). The 46% and 100% eluates were pooled and concentrated to 3 mL using Vivaspin 20 (MWCO 30 kDa) (VS2022, Sartorius) centrifugal ultrafiltration devices and then loaded at a flow rate of 1 mL per minute onto a HiLoad 16/60 Superdex75 pg (17-1068-01, Cytiva) previously equilibrated with SEC buffer (10 mM Hepes pH 7.5 (A1069, PanReac/AppiChem), 150 mM KCl (60130,Sima-Aldrich), 1 mM DTT (6908.4, Roth)). The bright green fractions, in the major peak running at the position expected for the monomeric form, were pooled together, concentrated, and the purity determined by SDS-PAGE.

### Nucleosome array acetylation *in vitro* and phase separation experiments

Two reaction conditions were prepared: −acetylation and +acetylation. For each condition, a master mix sufficient for seven subsequent reactions was assembled containing 14 µL chromatin (5 µM nucleosome concentration), 12.6 µL of 5X dilution buffer (125 mM Tris pH 7.5, 70 mM KCl, 25 mM DTT, BSA 0.5 mg/mL, 25% Glycerol), 7 µL of 5 µM p300 HAT-domain (produced by Protein Technologies, VBC) in 1X Dilution buffer, and 19.6 µL ddH_2_O. For the +acetylation condition, 6.3 µL of 10 mM acetyl-CoA (in ddH2O) was added, while the −acetylation condition received an equal volume of ddH_2_O. Reactions were incubated for 1 h at room temperature. The reactions were then quenched by addition of 3.5 µL 100 µM A-485 (in 1% DMSO) and samples were then incubated for 30 min at room temperature. 9 µL of each reaction was aliquoted into fresh tubes, nanobodies were added (1 µL of either 10 µM or 20 µM nanobody in 1X dilution buffer per tube), and control reactions received 1 µL of 1X dilution buffer. Samples were incubated for 30 min at room temperature, followed by addition of 10 µL of 2X Phase separation buffer (25 mM Tris pH 7.5, 300 mM KCl, 2 mM MgCl, BSA 0.1 mg/mL, Glycerol 5%) to each tube. Samples were loaded immediately into PEG coated 384-well plate with a glass bottom (coated as described in Gibson et al., 2019) and incubated for 1.5 h before imaging with a Zeiss LSM780 confocal microscope at room temperature. Remaining reaction material was kept for SDS-PAGE analysis and western blotting with acetylation specific antibody to assess for the acetylation reaction.

## Statistical analyses and data reporting

No sample sizes were predetermined using statistical methods.

## Data availability

Raw data are available and will be provided by the corresponding authors upon request. No restrictions apply to the availability of data provided within this manuscript.

## Use of large language models

ChatGPT (version 5.2, OpenAI) was used to assist in improving the grammar and editorial style of the manuscript text. No content was generated or modified relating to the scientific findings, data interpretation, or conclusions. All edits were reviewed and approved by the authors.

---

**Supplementary Figure 1.**
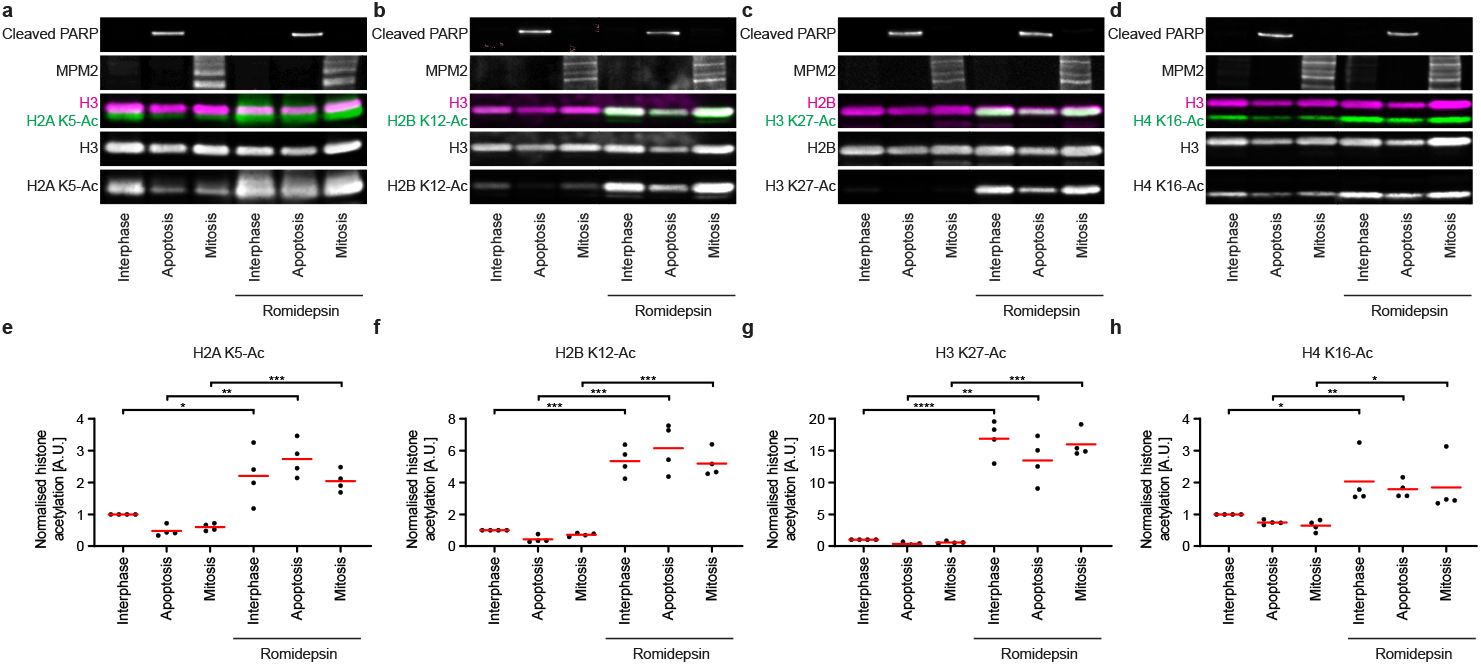
Romidepsin inhibits histone deacetylation without preventing apoptosis. **a-d**, Fluorescence-based western blot of histone acetylation levels of four core histones during apoptosis and mitosis, in the presence and absence of romidepsin. Cell lysates were harvested from control and romidepsin-treated interphase, apoptotic and mitotic cultures, and levels of histone acetylation were measured relative to core histones. Cleaved PARP and MPM2 were blotted to detect apoptotic and mitotic states, respectively, in both control and romidepsin-treated conditions. **e-h**, Quantification of histone acetylation levels, relative to core histone amounts in the same loading lane. *n*=4 for all conditions. Bars indicate mean; significance was tested by a parametric ratio paired t-test (H2A K5-Ac, *P*=0.0412 (interphase), *P*=0.0047 (apoptosis), *P*=0.0004 (mitosis); H2B K12-Ac, *P*=0.0003 (interphase), *P*=0.0005 (apoptosis), *P*=0.0009 (mitosis); H3 K27-Ac, *P*=<0.0001 (interphase), *P*=0.0028 (apoptosis), *P*=0.0004 (mitosis); H4 K16-Ac, *P*=0.0333 (interphase), *P*=0.0037 (apoptosis), *P*=0.0166 (mitosis)).

**Supplementary Figure 2.**
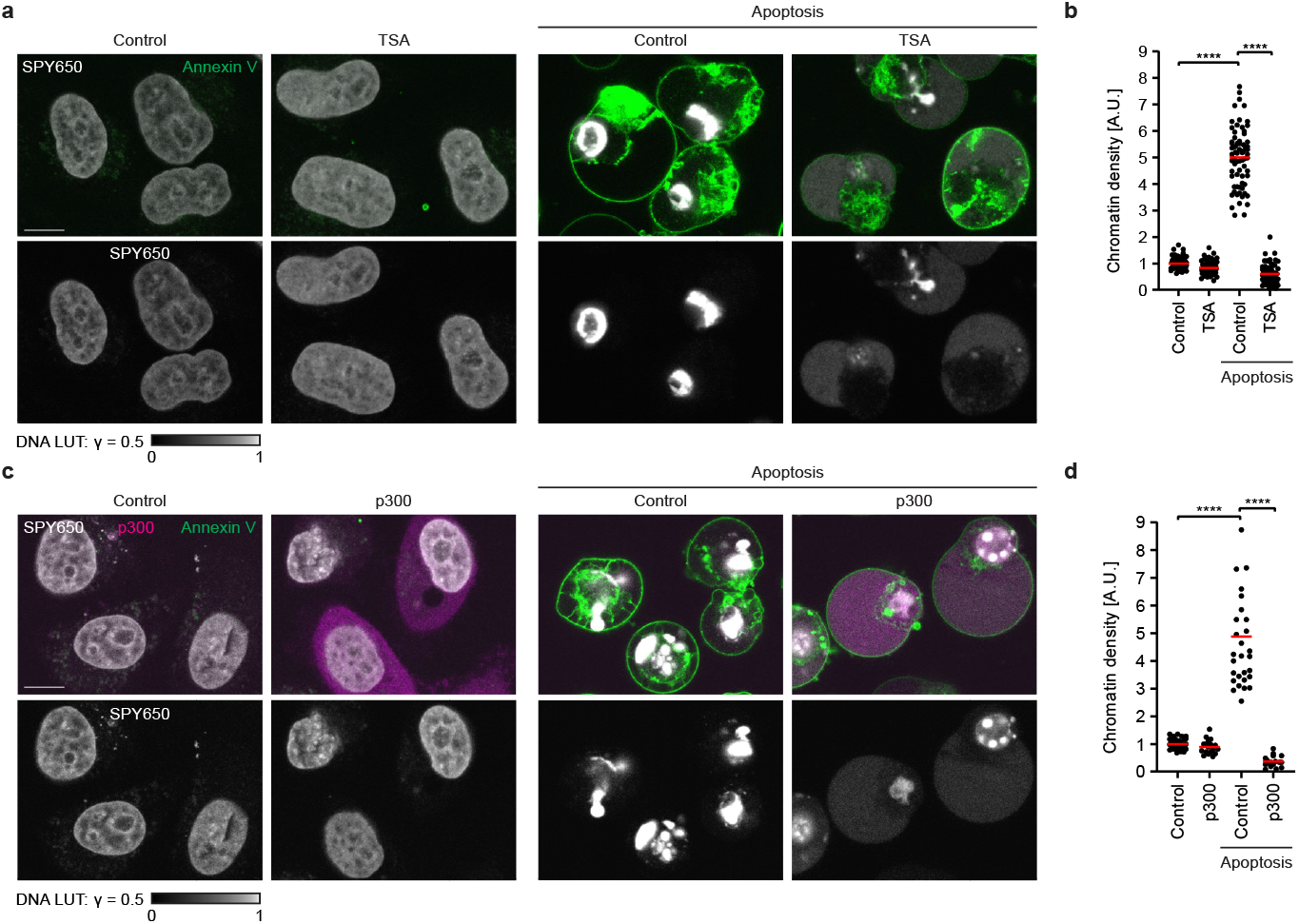
Diverse histone acetylation perturbations suppress apoptotic chromatin compaction. **a**, Imaging of chromatin density in the presence and absence of TSA in interphase and apoptosis. Chromatin was stained with the DNA dye SPY650, and Annexin V-488 was used to detect loss of membrane asymmetry in apoptosis. Scale bar 10 µm. **b**, Quantification of chromatin compaction. *n*=70 for interphase, *n*=71 for apoptosis (control); *n*=73 for interphase, *n*=68 for apoptosis (TSA). Bars indicate mean; significance was tested by a two-tailed Mann-Whitney test (control apoptosis vs control interphase, *P*=<0.0001; TSA-treated apoptosis vs control apoptosis, *P*=<0.0001). **c**, Imaging of chromatin density in the presence and absence of p300-GFP overexpression in interphase and apoptosis. Chromatin was stained with the DNA dye SPY650, and Annexin V-Pacific Blue was used to detect loss of membrane asymmetry in apoptosis. Scale bar 10 µm. **d, e**, Quantification of chromatin compaction. *n*=41 for interphase, *n*=30 for apoptosis (control); *n*=19 for interphase, *n*=18 for apoptosis (p300). Bars indicate mean; significance was tested by a two-tailed Mann-Whitney test (control apoptosis vs control interphase, *P*=<0.0001; p300 apoptosis vs control apoptosis, *P*=<0.0001).

**Supplementary Figure 3.**
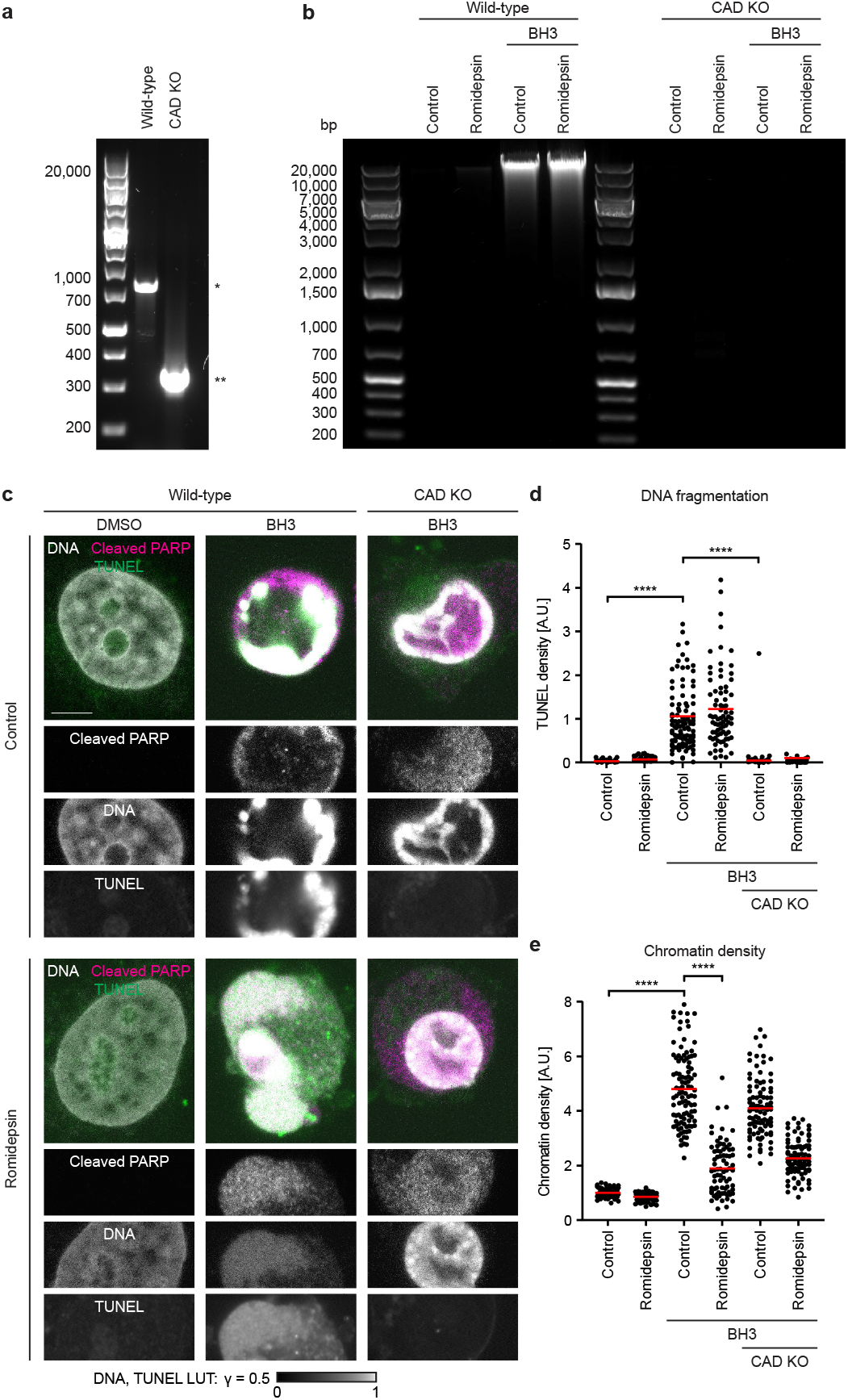
HeLa apoptotic DNA fragmentation fully depends on CAD and is unaffected by romidepsin treatment. **a**, Agarose gel electrophoresis after PCR amplification of the CAD genomic locus in wild-type and CAD KO HeLa cells. * indicates expected length of wild-type locus, ** indicates expected length of locus after CRISPR/Cas9-mediated partial deletion. **b**, Agarose gel electrophoresis of fragmented DNA isolated from wild-type and CAD KO cells in the presence and absence of romidepsin and BH3 mimetics. **c**, Imaging of fixed wild-type and CAD KO cells, stained for cleaved PARP and TUNEL signal. DNA was stained with SPY650. Scale bar 5 µm. **d**, Quantification of TUNEL signal normalised to chromatin density (**d**) and chromatin density (**e**) in fixed live and apoptotic wild-type and CAD KO cells, in the presence and absence of romidepsin. *n*=76 for live wild-type, *n*=103 for apoptotic wild-type, *n*=103 for apoptotic CAD KO (control); *n*=63 for live wild-type, *n*=77 for apoptotic wild-type, *n*=103 for apoptotic CAD KO (romidepsin). Bars indicate mean; significance was tested by a two-tailed Mann-Whitney test (apoptotic wild-type vs live wild-type, *P*=<0.0001 (TUNEL), *P*=<0.0001 (chromatin); apoptotic CAD KO vs apoptotic wild-type, *P*=<0.0001 (TUNEL), romidepsin-treated apoptotic wild-type vs control apoptotic wild-type, *P*=<0.0001 (chromatin)).

**Supplementary Figure 4.**
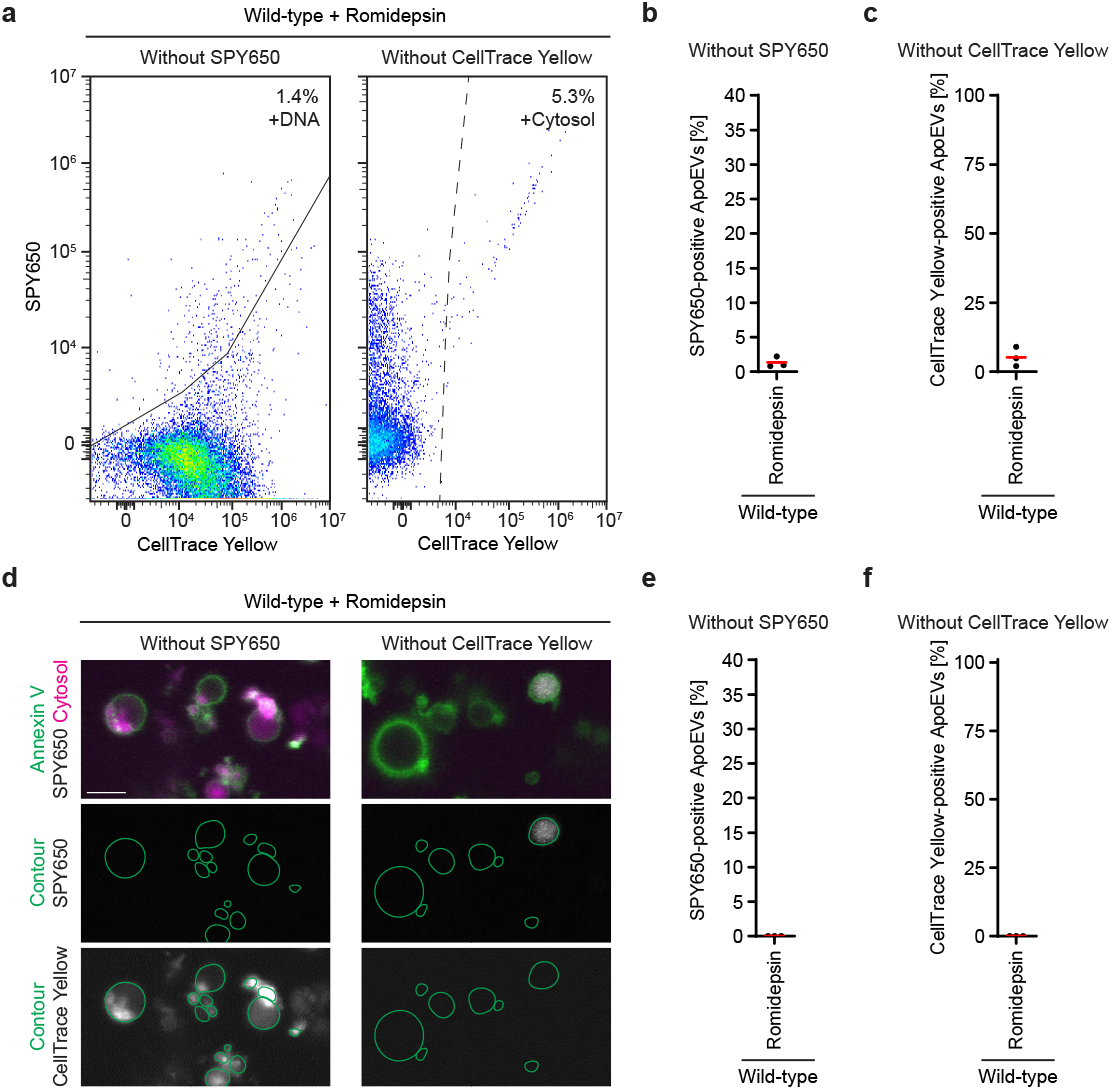
Characterisation of fluorescent background and defining positive populations for staining of ApoEVs. **a**, Flow cytometry plots of background ApoEV DNA and cytosol staining signal in romidepsin-treated apoptotic wild-type cells, under full-minus-one (FMO) staining. ApoEVs were identified based on size and Annexin V-488 staining, DNA was stained with SPY650 and cytosol with CellTrace Yellow. Solid line represents threshold for positive DNA staining, dotted line represents threshold for positive cytosolic staining. Thresholds were drawn manually, at the lowest point that resulted in exclusion of FMO ApoEVs. DNA percentages displayed as mean values across three biological replicates. **b, c**, Quantification of background ApoEV DNA (b) and cytosol (c) content, as a percentage of all ApoEVs. *n*=3 for all conditions. Bars indicate mean. **d**, Imaging of background DNA and cytosol staining within ApoEVs, under FMO staining. ApoEVs were stained with Annexin V-488, DNA with SPY650 and cytosol with CellTrace Yellow. Contours were drawn from the outer contour of Annexin V staining. Scale bar 5 µm. **e, f**, Quantification of imaged background ApoEV DNA (**e**) and cytosol (**f**) staining, as a percentage of all ApoEVs. *n*=3 for all conditions. Bars indicate mean.

**Supplementary Figure 5.**
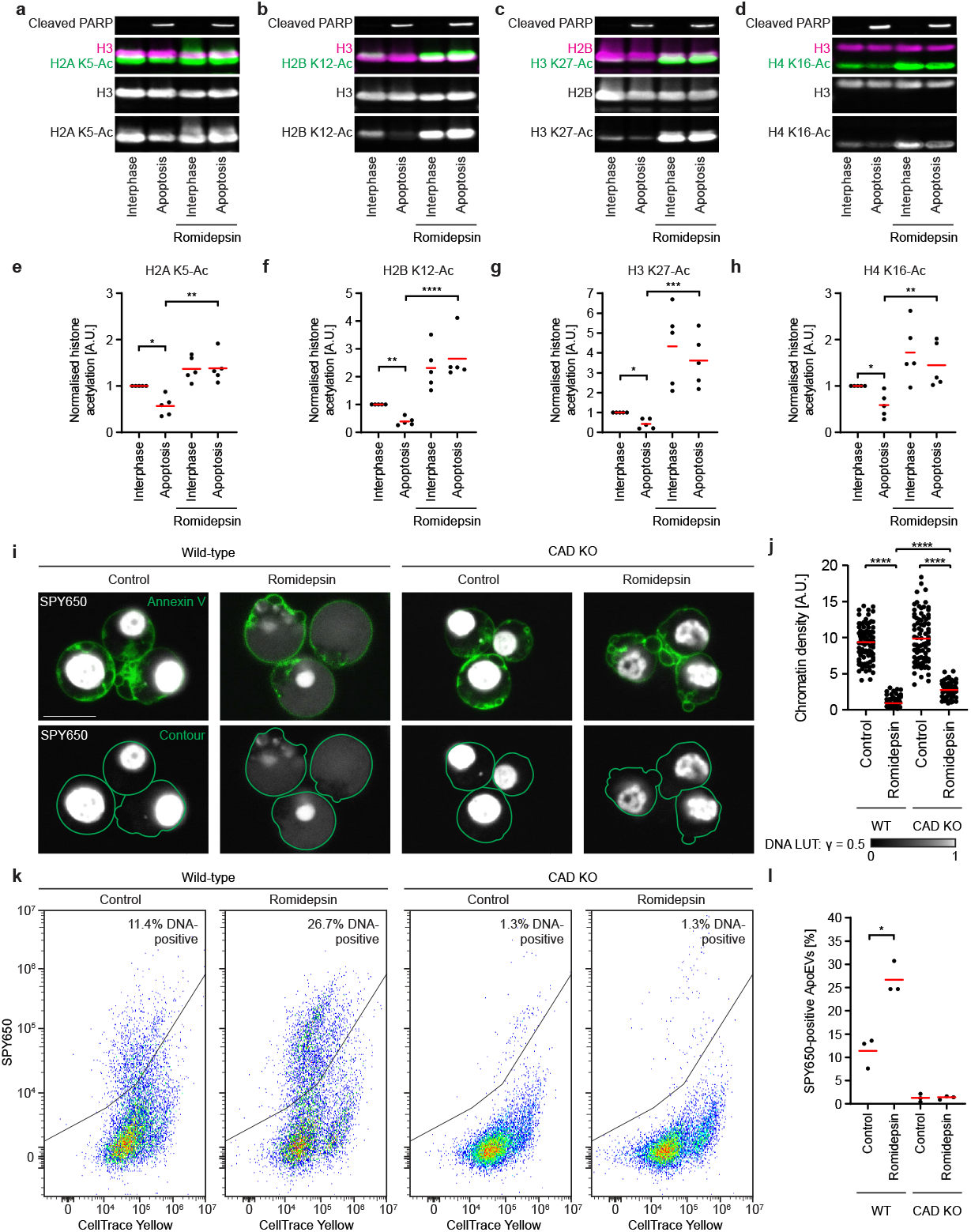
Apoptotic histone deacetylation is required to compact and sequester fragmented DNA in the human Jurkat cell line. **a-d**, Fluorescence-based western blot of histone acetylation levels of four core histones during apoptosis in Jurkat cells, in the presence and absence of romidepsin. Jurkat cell lysates were harvested from control and romidepsin-treated interphase and apoptotic cultures, and levels of histone acetylation were measured relative to core histones. Cleaved PARP was blotted to detect an apoptotic state in both control and romidepsin-treated conditions. **e-h**, Quantification of Jurkat cell histone acetylation levels, relative to core histone amounts in the same loading lane. *n*=5 for all conditions. Bars indicate mean; significance was tested by a parametric ratio paired t-test (H2A K5-Ac, *P*=0.0215 (control apoptosis vs control interphase), *P*=0.0032 (romidepsin-treated apoptosis vs control apoptosis); H2B K12-Ac, *P*=0.0032 (control apoptosis vs control interphase), *P*=<0.0001 (romidepsin-treated apoptosis vs control apoptosis); H3 K27-Ac, *P*=0.0235 (control apoptosis vs control interphase), *P*=0.0002 (romidepsin-treated apoptosis vs control apoptosis); H4 K16-Ac, *P*=0.0452 (control apoptosis vs control interphase), *P*=0.0069 (romidepsin-treated apoptosis vs control apoptosis)). **i**, Imaging of chromatin density in apoptotic wild-type and CAD KO Jurkat cells in the presence and absence of active histone deacetylases. Chromatin was stained with the DNA dye SPY650, and Annexin V-488 was used to detect loss of membrane asymmetry in apoptosis. Contours were drawn from the outer contour of Annexin V staining. Scale bar 10 µm. **b**, Quantification of chromatin compaction in apoptotic Jurkat cells. *n*=94 for wild-type control apoptosis, *n*=93 for wild-type romidepsin-treated apoptosis, *n*=86 for CAD KO control apoptosis, *n*=73 for CAD KO romidepsin-treated apoptosis. Bars indicate mean; significance was tested by a two-tailed Mann-Whitney test (romidepsin-treated wild-type apoptosis vs control wild-type apoptosis, *P*=<0.0001; romidepsin-treated CAD KO apoptosis vs control CAD KO apoptosis, *P*=<0.0001; romidepsin-treated CAD KO apoptosis vs romidepsin-treated wild-type apoptosis, *P*=<0.0001). **k**, Flow cytometry plots of DNA content in ApoEVs from apoptotic wild-type and CAD KO Jurkat cells, in the presence and absence of romidepsin. ApoEVs were identified based on size and Annexin V-Pacific Blue staining, DNA was stained with SPY650. Positive populations were defined by unstained controls. DNA percentages displayed as mean values across three biological replicates. **l**, Quantification of Jurkat ApoEV DNA content, as a percentage of all ApoEVs. *n*=3 for all conditions. Bars indicate mean; significance was tested by a parametric paired t-test (DNA content in romidepsin-treated wild-type cells, *P*=0.0186).

**Supplementary Figure 6.**
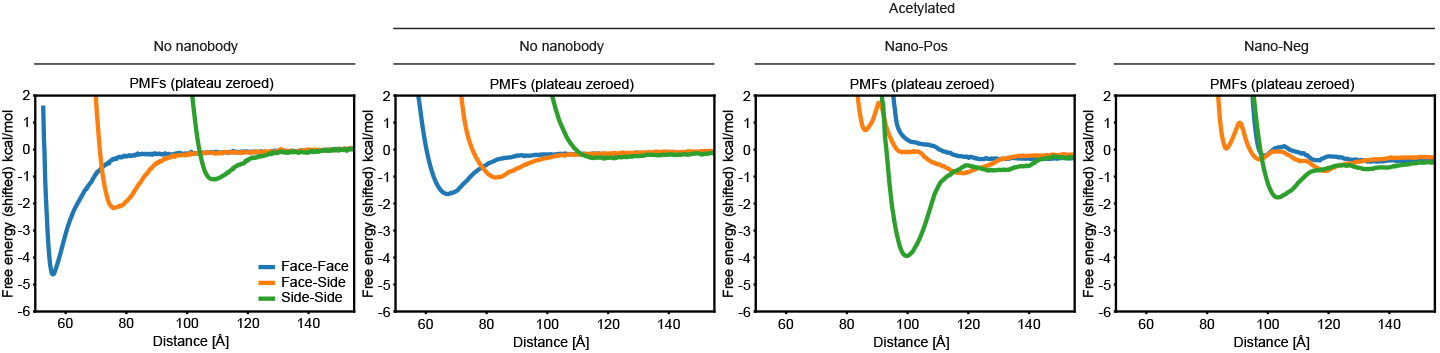
Potential-of-mean-force calculations of control nucleosome arrays and acetylated nucleosome arrays in the presence of Nano-Pos and Nano-Pos. Simulation time = 400 ns for all conditions.

**Supplementary Table 1.**
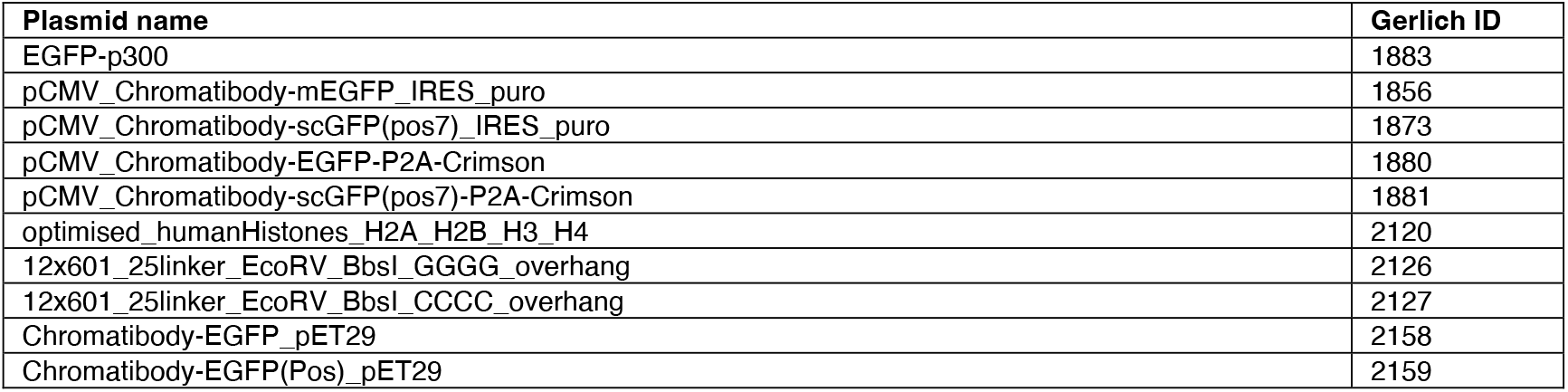
A list of plasmids used in this study, and accompanying Gerlich lab IDs.

**Supplementary Table 2.** A list of antibodies used in this study, including supplier, catalogue number, lot number and accompanying Gerlich lab IDs.

| Antibody | Gerlich ID | Supplier | Catalogue # | Lot # |
| --- | --- | --- | --- | --- |
| Rabbit anti-Histone H2A (acetyl K5) | 468 | Abcam | ab45152 | 1038245-8 |
| Rabbit anti-Acetyl Histone H2B (Lys12) | 496 | CST | 5410S | 1 |
| Rabbit anti-Acetyl Histone H3 (Lys27) (D5E4) | 357 | CST | 8173S | 9 |
| Rabbit anti-Histone H4k16 (acetyl K16) | 372 | Abcam | ab109463 | 3357884-3 |
| Rabbit anti-H4K20Ac | 375 | Invitrogen | MA5-33387 | VG3018089 |
| Mouse anti-Histone H3 | 206 | Active motif | 39763 | 25031213 |
| Mouse anti-Histone H2B - ChIP Grade | 327 | Abcam | ab52484 | 1078693-1 |
| Rabbit anti-Cleaved PARP (Asp214) (D64E10) | 476 | CST | 5625S | 18 |
| Mouse anti-Phospho-Ser/Thr-Pro MPM-2 | 388 | Sigma | 05-368 | 3485885 |
| Goat anti-rabbit IRDye 800CW | 165 | Li-COR | 926-32211 | D50311-04 |
| Donkey anti-mouse IRDye 680RD | 169 | Li-COR | 926-68072 | D41217-05 |
| Goat anti-rabbit IRDye 680LT | 164 | Li-COR | 926-68021 | C30820-02 |
| Donkey anti-mouse IRDye 800CW | 170 | Li-COR | 926-32212 | C31024-04 |
| Donkey anti-rabbit IgG Alexa Fluor™ 594 | 056 | Thermo | A-21207 | 1744751 |

**Supplementary Table 3.** A list of cell lines used in this study, and accompanying Gerlich lab IDs.

| Cell line name | Gerlich ID |
| --- | --- |
| HeLa Kyoto wild-type | 0001 |
| HeLa Kyoto CAD KO | 2516 |
| Jurkat E6-1 wild-type | 2301 |
| Jurkat E6-1 CAD KO | 2421 |

